# Fish microbiomes 101: disentangling the rules governing marine fish mucosal microbiomes across 101 species

**DOI:** 10.1101/2022.03.07.483203

**Authors:** Jeremiah Minich, Andreas Härer, Joseph Vechinski, Benjamin W Frable, Zachary Skelton, Emily Kunselman, Mike Shane, Daniela Perry, Antonio Gonzalez, Daniel McDonald, Rob Knight, Todd P. Michael, Eric E Allen

## Abstract

Fish are the most diverse and widely distributed vertebrates, yet little is known about the microbial ecology of fishes nor the biological and environmental factors that influence the fish microbiome. The microbiota from 101 species of Southern California marine fishes, spanning 22 orders, 55 families, and 83 genera representing ~25% of local marine fish diversity, was analyzed to identify patterns that explain microbial diversity patterns in a geographical subset of marine fish biodiversity. We compared fish microbiomes (gill, skin, midgut, and hindgut) using alpha, beta, and gamma diversity along with establishing a novel method to estimate microbial biomass (Qiime2 plugin katharoseq). For oceanic fishes from the neritic zone, host size and distance from shore were negatively associated with microbial biomass densities and diversity in the gills. Body site was the strongest driver for beta diversity with strong evidence of phylosymbiosis observed across the gill, skin, and hindgut, but not midgut. The majority of microbes from all fish body sites were of unknown origin but overall sea water generally contributes more microbes to fish mucus compared to marine sediment. In a meta-analysis of vertebrate hindguts (569 species), mammals had the highest gamma diversity when controlling for host species number while fishes had the highest percent of unique microbial taxa (92%). In fishes, the midgut, gill, and skin contains the majority of microbial diversity which collectively can be 5.5 times higher than the hindgut. The composite dataset will be useful to vertebrate microbiome researchers and fish biologists interested in microbial ecology with applications in aquaculture and fisheries management.

## INTRODUCTION

The earth is estimated to contain around 1 trillion microbes ^1^ and while several efforts have sought to describe these communities as they relate to their environmental biomes ^2,3^, few studies have focused on non-human vertebrates ^4,5 6^. The host-associated gut microbiome in vertebrates is shaped by a variety of biological factors including phylogeny, diet, and age, along with environmental factors such as geography, habitat, and climate, whereas less is known about other body sites ^4,5,7–10^. Of the large meta-analyses that have sought to evaluate vertebrate host microbial diversity, most focus exclusively on hindgut or stool from terrestrial animals from captive (zoo) environments. Fishes, despite being the most phylogenetically diverse vertebrates, are severely underrepresented in these studies ^4,9^. The underrepresentation of fishes in these datasets is a critical concern because many theories have arisen from these studies, including phylosymbiosis, contributions of diet driving community assemblies, while body sites outside the gut are ignored and aquatic animals are insufficiently sampled to establish the generality of conclusions.

Fishes include several broad classes collectively representing the largest diversity of species within the vertebrate classes: Agnatha (jawless), Chondrichthyes (cartilaginous), Sarcopterygii (lobe-finned), and Osteichthyes (bony). This study aimed to answer three main questions to understand the ecological and biological drivers of the fish microbiome: 1) what are the primary factors that influence host-associated microbial communities for marine fishes (e.g. body site location, habitat, trophic level, swimming method, phylogeny, etc.); 2) where might these microbes originate (sea water, sediment, host, etc); and 3) is microbial diversity greater in fishes compared to other vertebrates considering fishes are evolutionarily more ancient. We sampled and analyzed the microbiota from the four primary fish mucosal body sites (gill, skin, midgut, and hindgut) for 101 species (28 orders, 55 families, and 83 genera) of marine fishes from Southern California (SoCal; Eastern Pacific Ocean ‘EPO’), which represent approximately 25% of the local marine fish diversity, to quantify impacts of host phylogeny, trophic level, habitat type, swim performance, and body site. We also included gill samples from 17 species of fishes from the Atlantic, including two species also in the EPO dataset, bringing the total to 30 orders, 61 families, 96 genera, and 116 species.

## RESULTS

### Sampling and microbial biomass estimation

From March 2018 through September of 2020 fish were collected during both directed sampling efforts along with passive sampling, namely through donations from recreational anglers. Fish were caught from a diverse range of nearshore and offshore habitats along with a range of depths (0 to 500 m) in the EPO and Western Atlantic (Figure 1a-b). Standard biometric measures were taken for all fish. For the 101 species from the EPO, a total of four body sites (gill, skin, midgut, and hindgut) were sampled for microbiome analysis whereas only gill samples were processed for the Atlantic species subset (15 additional unique species) (Figure 1c). A final table with all of the fish used in the study alphabetically sorted by order, family, and then species name along with corresponding pictures of the fish and a gill sample can be found in Supplemental File 1.

**Figure 1.**
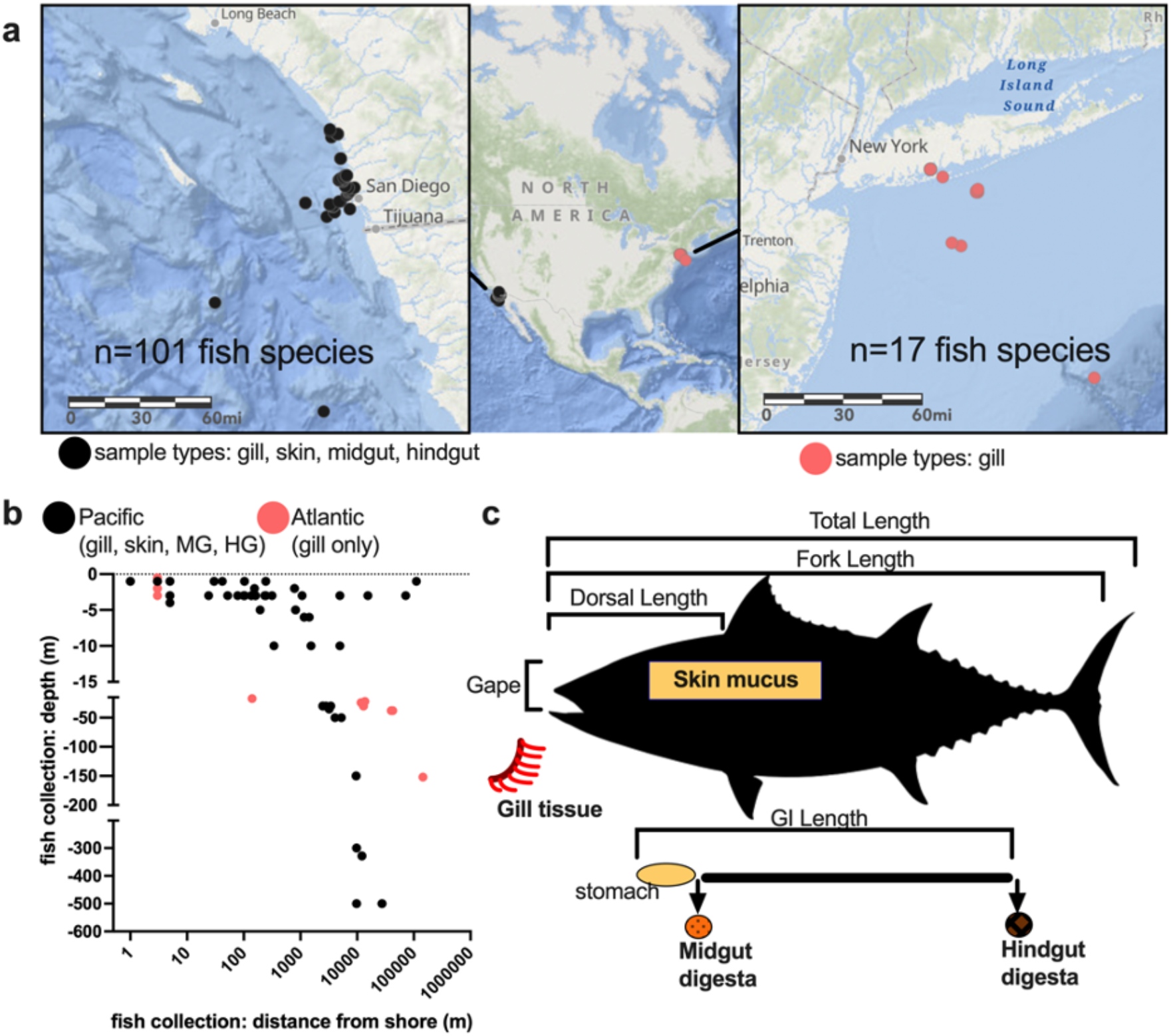
Sampling design of 116 species of marine fish. A) Using ArcGIS to depict the general area from which fish were sampled: black dots indicate the locations of the 101 unique species of marine fish sampled from the California Current Ecosystem in the Eastern Pacific Ocean primarily in the waters of San Diego CA. Red circles depict the locations of an additional 17 species of fish (15 unique species with 2 species duplicates) collected from the Western Atlantic primarily in the waters of New York. When multiple species of fish were caught in the same location, a single circle is used to indicate the location. B) Fish were sampled across a gradient of depth and distances from shore. C) Biometric measurements taken for nearly all fish include total length, fork length, mass, gape, and GI length. Various ratios from these lengths were also calculated. Microbiome samples from the gill were primarily whole tissue specimens from the entire left second gill arch or a section of the top middle and bottom of the entire filament. Skin mucus samples were taken by scraping using a razor blade. Midgut digesta material was collected from directly posterior of the stomach or if stomach was absent, the beginning of the GI tract. Hindgut digesta samples were taken from near the anus.

### Microbial biomass estimation

Katharoseq was applied to determine the limit of detection of the assay, whereby the total post-deblur read counts of the positive DNA extraction controls were compared to the relative abundance of the known targets (Figure 2) ^11^. The 0.9 threshold was applied and the read number at y=0.9 was determined to be 1150 reads (Figure 2a). Thus, any samples with less than 1150 reads were excluded from the fish microbiome project (FMP) analysis. Since a subset of the standards had known cell concentrations, we then determined the limit of detection of the assay based on cell counts (in addition to the read counts). At y=0.9, the number of input cells to the DNA extraction was estimated to be 15.95 (Figure 2b). Since the positive controls (*Bacillus subtilis* and *Paracoccus spp*.) used in extraction generally have a high 16S copy number (~10 rRNA copies per genome), we estimated the limit of detection of the assay to be between ~16-160 microbial cells. Next we log transformed the reads and known cell quantities to generate a model that enables one to predict cell counts from read counts (P<0.0001, R2=0.8668, m=3.497) (Figure 2c). Samples that did not have at least 1150 reads were excluded per above. We used this equation to estimate the microbial biomass for each sample in the FMP dataset and then extrapolated for the total volume (ul) in the extraction and finally normalized by the mass (g) of tissue used into the extraction to get a final value of microbial cells per g of tissue. The actual distribution of taxonomy (target controls shown) within the positive titrations is displayed (Figure 2d). Using the 1150 read cutoff, we excluded any samples in the FMP dataset with less than 1150 reads which overall yielded a high success rate across the various sample types (Figure 2e).

**Figure 2.**
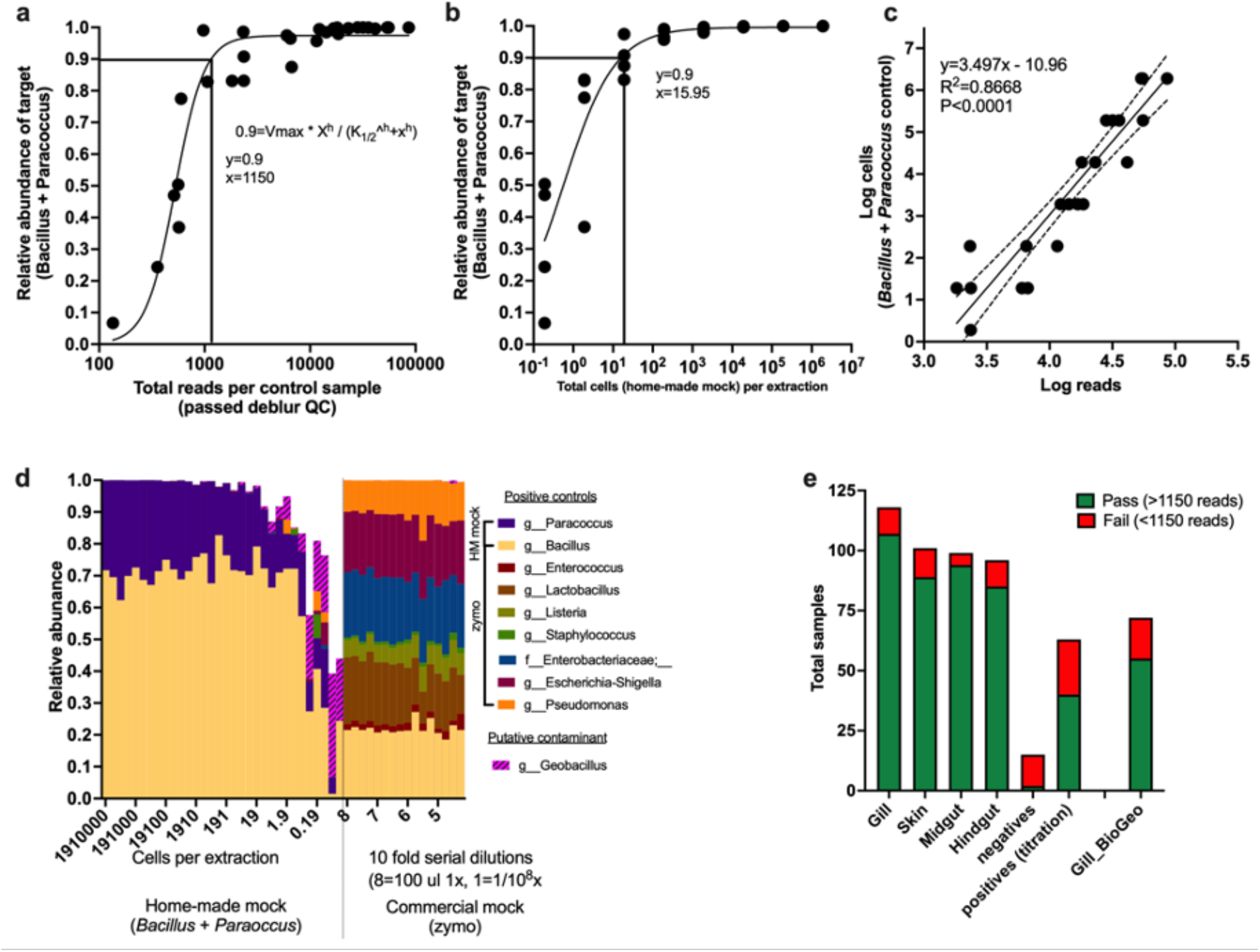
Limit of detection, sample exclusion, and microbial biomass estimation for FMP101 dataset. a) application of Katharoseq formula to calculate limit of detection of microbiome plates using the *Bacillus* / *Paracoccus* mock community (1150 reads at 90%). b) limit of detection based on cell counts of *Bacillus* / *Paracoccus* mock community (~16 cells into extraction at 90%). c) Model fit of the log(sequencing read counts) of positive extraction controls vs. the log cell counts of those positive extraction controls (empirically determined using plate counts. The linear regression of the line is indicative of the quality of method to estimate microbial biomass from sequencing read counts. This method is similar to a qPCR curve where the log (Ct) would be equivalent to the log(read counts). This equation is then used to estimate the number of ‘microbial density’ of the existing samples which is then further normalized by the volume of the DNA extraction, biomass of material going into the extraction and finally normalized to at estimated microbial cells per gram of tissue. d) community analysis comparison and validation of compositionality of controls of two sets of mock community controls (section 1 = *Bacillus* / *Paracoccus* mock community; section 2 = zymo mock community). Putative contaminant g__*Geobacillus* identified (present in 93% of negatives and higher relative abundance as compared to positives and samples). e) number of samples successful across the four body sites collected from the broad fish microbiome dataset.

Since using a non-rarified (or raw count) dataset is somewhat contentious in the field due to the argument of not knowing absolute abundances, and although we showed that estimating actual microbial abundances is feasible, we performed additional beta diversity testing to determine how these two strategies may impact interpretation of results (Supplemental Fig. 2). Specifically, we either rarified all samples at 1150 reads (excluding samples with less than 1150 reads) or we simply excluded samples with less than 1150 reads, keeping the raw counts (non-rarified). We also tested the effects of removing chloroplast reads from the dataset, which is a common contaminant in aquatic microbiome datasets. When comparing overall trends in the dataset, the order of significant drivers of the microbiome was generally conserved and not altered when comparing both processing methods. On a per factor basis, for Unweighted UniFrac specifically, certain factors had a slightly higher F-stat for the rarified version versus the non-rarified, but the differences were minimal (Supplemental Fig. 2a). The decision to remove or retain chloroplasts did not change the effect size or F-stat for Unweighted UniFrac. For Weighted Unifrac the decision to rarefy or not was even less drastic with all orders of important factors remaining unchanged. Moreover, removing chloroplasts generally did not influence the order except for a flip between habitat_depth_level1 and substrata_group, whereby the chloroplasts would have a stronger influence of community differentiation when comparing shallow (neritic), midwater mesopelagic, and bathypelagic zones. Therefore, we proceeded by using the non-rarified dataset with samples having less than 1150 reads and removed chloroplasts. The final FMP dataset includes a total of 373 successful samples including 107 gill samples, 89 skin, 94 midgut, and 85 hindgut (Figure 2e).

### Factors influencing alpha diversity of the fish microbiome

Next we evaluated the primary factors that influence the microbial communities in the marine fish mucosal samples. First we compared the alpha diversity metrics (Chao1, Faith’s Phylogenetic Diversity, and Shannon) and microbial biomass (estimated microbial cells per g tissue) across sample type (body site) from all fishes to determine if certain body sites had unique microbial signatures. For all three alpha diversity metrics: Chao1 (Figure 3a, P<0.0001 KW=56.58), Faith’s PD (Figure 3b, P<0.0001, KW=45.95), and Shannon (Figure 3c, P<0.0001, KW=47.43), there were significant differences across body sites. For all alpha diversity metrics, the midgut samples had the highest diversity compared to other body sites (gill, skin, hindgut), while skin had higher diversity than gill. For Shannon diversity only, skin was higher than hindgut (Figure 3c). When comparing microbial biomass, there were no significant differences across body sites (Figure 3d), although the range of biomass was substantial (over 6 orders of magnitude).

**Figure 3.**
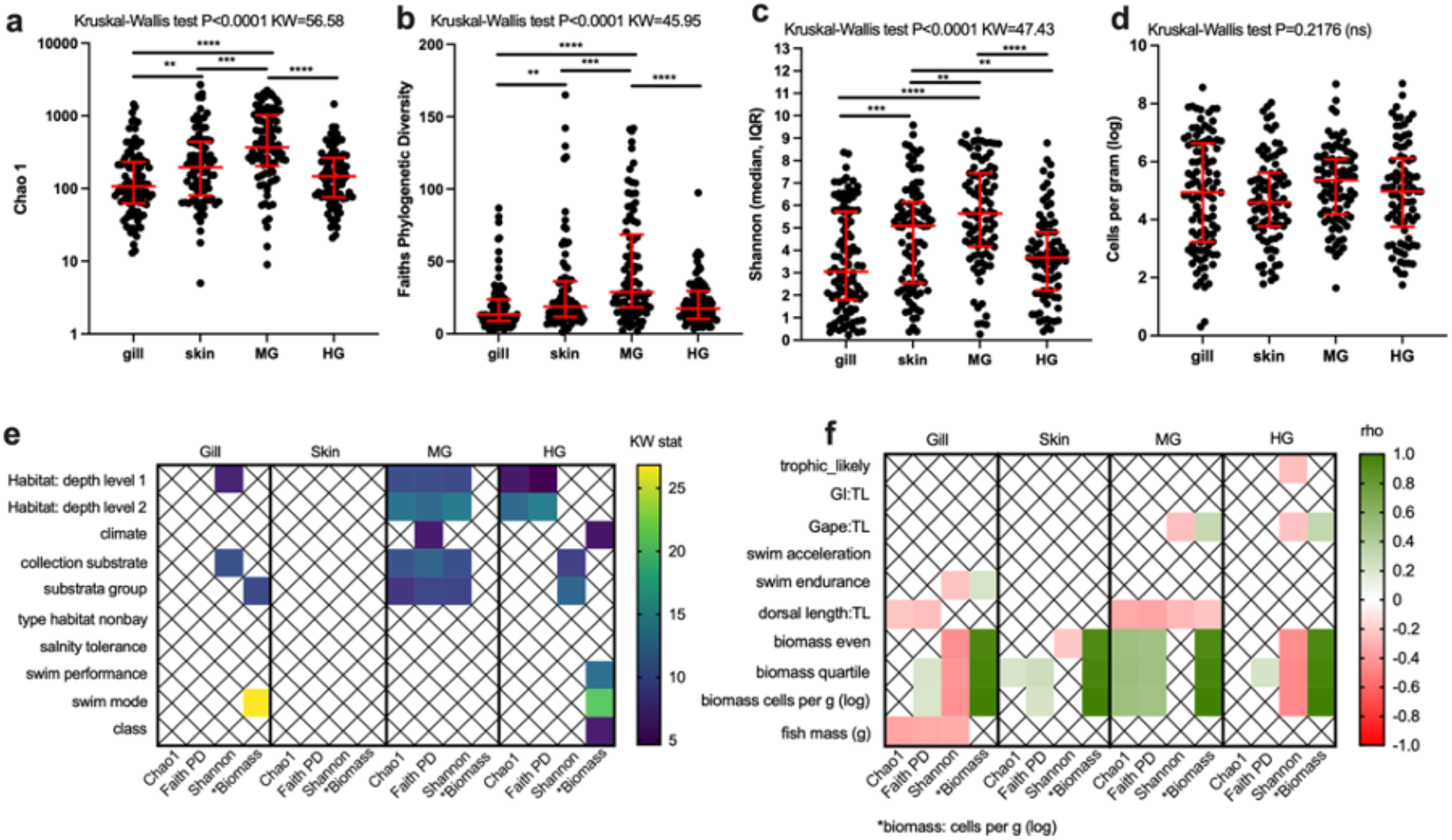
Alpha diversity and biomass comparisons across ecological and biological gradients in marine fish. Comparison of microbial diversity a) “Chao1”, b) “Faith’s Phylogenetic Diversity, c) “Shannon”, or d) microbial biomass across body site (gill, skin, midgut, and hindgut). Statistical differences determined using non-parametric testing Kruskal-Wallis test with 0.05 FDR Benjamini-Hochberg. Further testing computed for each unique body site for a variety of biological and ecological metadata categories. Metadata which is e) categorical is tested using Kruskal-Wallis f) whereas numeric metadata tested using Spearman correlation. Only significant associations are represented in e (Krukal-Wallis P<0.05) and f (Spearman P<0.05).

Various life history metrics were categorized for all fishes (Figure 3; Supplemental Table 1. FMP.alpha). For categorical variables, a Kruskall-Wallis test was used to compare the three alpha diversity metrics and biomass (log microbial cells per gram of fish tissue). For gill samples, Shannon diversity was influenced by habitat depth level 1 and collection substrate, whereas biomass was influenced by substrata group and swim mode (Figure 3e). Variation in skin samples was not explained by any of the categorical metadata. The midgut had the highest amount of differences in alpha diversity explained by the metadata categories. For habitat level 1, habitat level 2, collection substrate and substrata group differences were observed for Chao1, Faith’s PD, and Shannon diversity. In addition, the climate category that refers to the approximate latitudinal range and water temperature regime the fishes reside (temperate, subtropical, or tropical), biomass differed (Figure 3e). For the hindgut samples, habitat level 1 and habitat level 2 influenced the Chao1 and Faith’s PD. Shannon diversity was influenced by collection substrate and substrata group. Lastly, biomass was associated with climate, swim performance and swim mode (Figure 3e).

Next, we assessed the continuous or numeric values using Spearman correlation across the alpha diversity and biomass metrics for each body site. Some of these metrics are ratios, e.g. RIL (relative intestinal length) = total gut length/total fish length. For the trophic associated metadata categories, only the hindgut Shannon diversity was significant, and was negatively associated with high trophic level (Figure 3f). For the swim associated metrics, acceleration was not associated with any metric across the body sites, whereas swim endurance was positively associated with microbial biomass in the gill and negatively associated with Shannon diversity in the gill (Figure 3f). The dorsal length:total length ratio was negatively associated with Chao1 and Faith’s PD in the gill and negatively associated with all metrics in the midgut. This would suggest that faster swimming fishes have lower gill microbiome diversity and lower midgut microbiome diversity. For the biomass measurements, Shannon diversity in the gill and hindgut were negatively associated with microbial biomass. Faith’s PD was positively associated with microbial biomass in the midgut and partially associated in gill, skin, and hindgut. Chao 1 was positively associated with microbial biomass in midgut and partially in skin and hindgut (Figure 3f). Lastly, when comparing the total mass of the fishes against microbial biomass, there were no significant associations in the skin, midgut, or hindgut, whereas all of the gill microbiome alpha diversity metrics were negatively associated with mass of fishes (Figure 3f).

A subset of fish samples from the neritic zone (0-200 m) and collected exclusively from the ocean (excluding samples collected from bays or estuaries) was analyzed to further evaluate microbiome associations on the fish gill. We did this to control for and reduce effects from environmental noise associated with bays such as salinity gradients and tidal flow along with temperature for deep sea fish. Specifically we tested whether the mass of the fish or the distance from shore from which the fish was caught predicted microbiome characteristics in the fish gill. Since both fish mass and distance from shore were positively correlated (Spearman P<0.001, rho = 0.43), interpretation of results should be with caution as we were not able to tease apart these confounding variables (Figure 4a). The gill microbial biomass differed across habitats from which the fish were caught (Kruskal-Wallis P=0.0144, KW=16.31) with pelagic fish having lower microbial biomass in the gill compared to fish collected from the intertidal and subtidal zones (P<0.05) (Figure 4b). This result led us to test if either fish mass or the distance from shore had an impact on microbial biomass or diversity as intertidal and subtidal fish are close to shore whereas pelagic primarily live offshore. Fish mass (Figure 4c) and distance from shore (Figure 4d) were both negatively associated with gill microbial biomass. In addition, fish mass and distance from shore were negatively associated with Chao1 (Figure 4e-f) and Faith’s PD (Figure 4f-g). Overall for oceanic fish living in the neritic zone, we observed that offshore fishes such as pelagics along with larger fishes have lower microbial biomass density and diversity in the gills as compared to small fish living closer to shore such as fish from the intertidal and subtidal environments.

**Figure 4.**
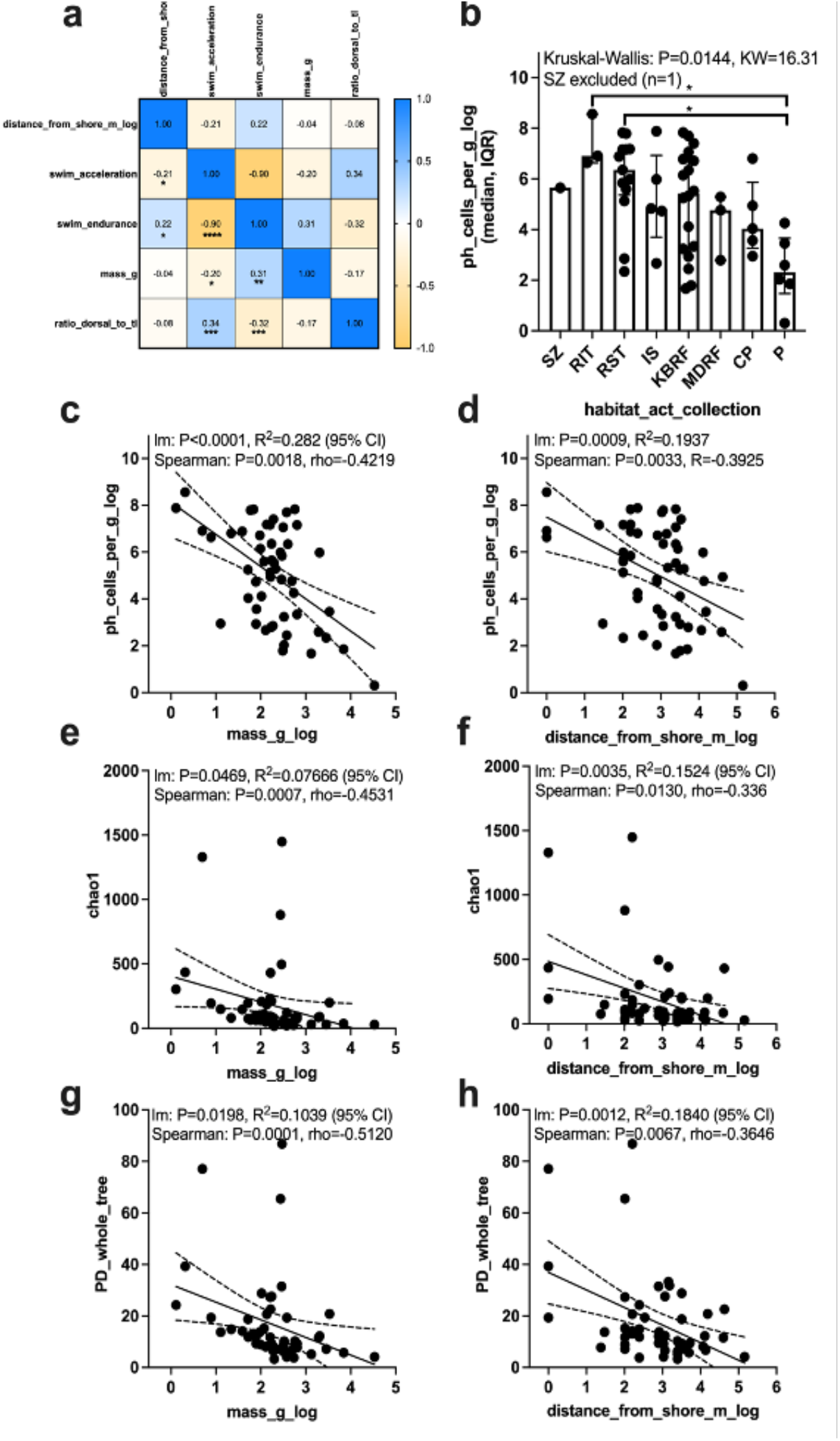
Associations between fish mass and collection location as measured by distance from shore with fish gill microbial biomass and alpha diversity. Subset of fish from EPO and Atlantic (n=54) collected from ocean (excludes bay and estuary samples) and from the neritic zone (<200 m depth). A) Correlation matrix between sample metadata where values are rho and significance indicated by * P<0.05, ** P<0.01, *** P<0.001, **** P<0.0001 (Spearman correlation). B) Comparison of gill microbial biomass (log cells per gram) across habitat types from which the fish were collected. C) Comparison of fish mass and D) distance from shore with gill microbial biomass. E) Comparison of fish mass and F) distance from shore with alpha diversity metrics (Chao1). G) Comparison of fish mass and H) distance from shore with alpha diversity metric: Faith’s PD.

### Factors influencing Beta diversity in the fish microbiome

We generally assessed the same biological and life history traits of the fish species for microbial beta diversity (Figure 5; Supplemental Table 2. FMP.beta). First, we compared all samples together (n=373) for Unweighted and Weighted normalized Unifrac (Figure 5a). Sample type (gill, skin, midgut, hindgut) for both metrics were the primary predictor. Microbial biomass, habitat depth (shallow ‘neritic’, mid ‘mesopelagic’, and deep ‘greater than 1000 m’), and substrata group (refers to benthic substrate type such as soft bottom, rocky reef, deep water, pelagic, etc.) were also large predictors. Since the sample type was the biggest driver, we subsampled each body site and analyzed again. Gill samples were associated with water column depth (habitat level), benthic substrate material, and microbial biomass (Figure 5a) for Unweighted Unifrac. There were generally fewer associations across body sites for Weighted Unifrac, but benthic substrate type was also associated with gill and hindgut communities (Figure 5a). Trophic level did not predict any beta diversity metrics for individual body sitesl. However, lower trophic level fishes had a higher similarity between midgut and hindgut, whereas more carnivorous fishes tended to have higher variation between midgut and hindgut (Figure 5b). Higher trophic fishes had more differentiation between the midgut and hindgut. Although lower trophic fishes had a higher GI length to body length ratio, this suggested additional factors may be a stronger influence than gut length.

**Figure 5.**
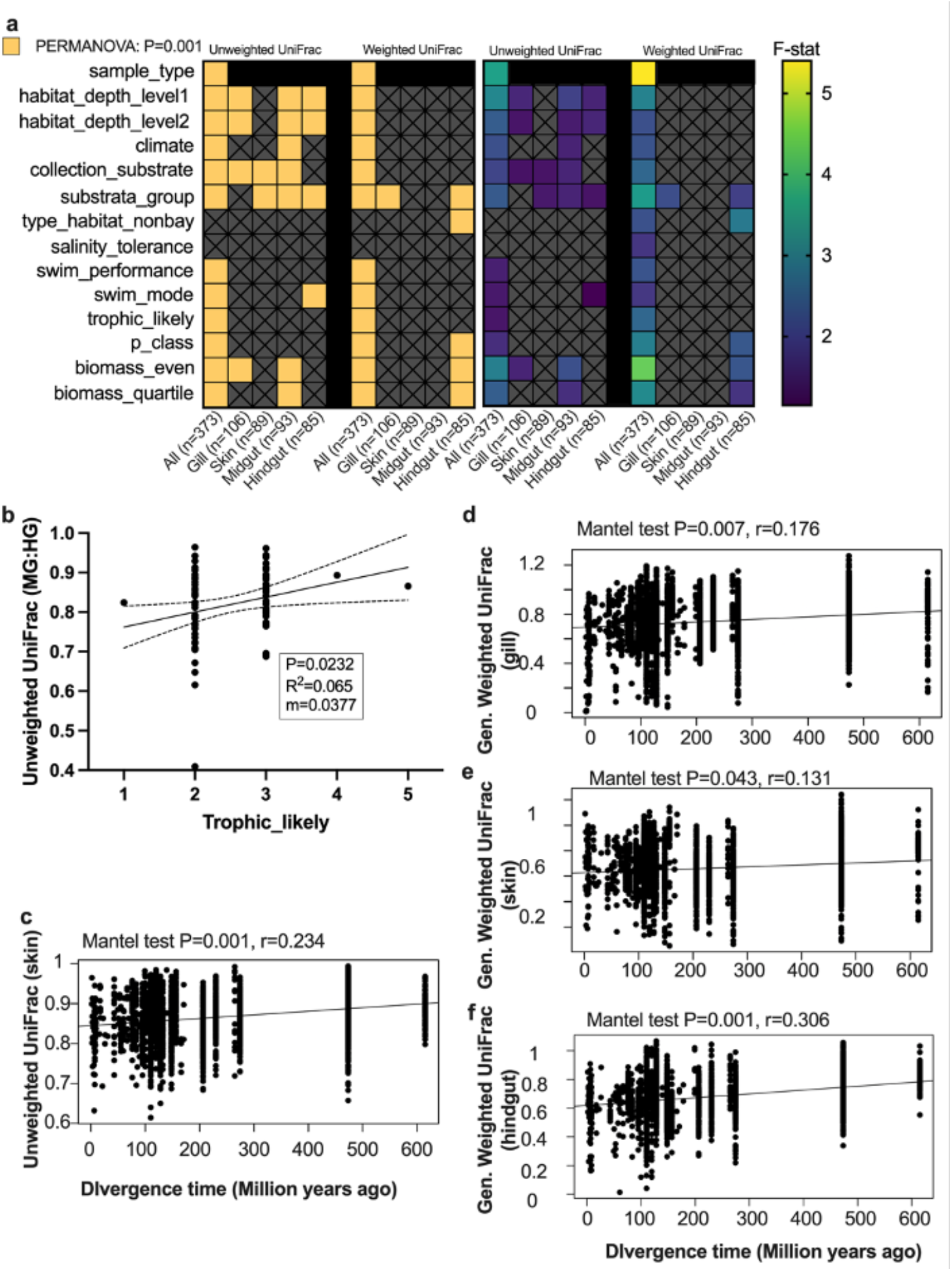
Biological and life history drivers of mucosal microbiomes in diverse sampling of marine fish from Southern California. a) Multivariate analysis of biological and life history parameters evaluated using Unweighted and Weighted normalized UniFrac distances. Statistical significance (PERMANOVA P=0.001) indicated by yellow blocks (left) and effect size (right). All samples compared together (all) along with individual sample types (gill, skin, midgut, hindgut). b) Impact of trophic level on similarity between midgut and hindgut (within a species) (linear model). Effect of evolutionary distance (similarity) of all fish compared to c) skin unweighted Unifrac distance, d) gill generalized weighted Unifrac distance, e) skin generalized weighted Unifrac distance, and f) hindgut generalized weighted Unifrac distance. Comparisons performed using Mantel test. Divergence time between fish species calculated using timetree.org.

### Evidence for phylosymbiosis across multiple body sites

Phylosymbiosis, the association of microbes with the phylogeny of the host, in vertebrates has been shown to occur in the hindgut ^12^ and skin ^13^ of mammals, but no studies have tested this hypothesis in fishes across multiple body sites. We evaluated if the evolutionary distance between fish species was associated with microbiome similarity across four body sites (gill, skin, midgut, and hindgut) when evaluated independently and measured by both Unweighted and Weighted normalized UniFrac. Using timetree.org we created a tree (Supplemental Fig. 3) to estimate phylogenetic distances between fishes that were compared to microbiome distances for each body site independently. When comparing microbiome distance based on unweighted Unifrac to evolutionary distances in fishes, only the skin was associated (p=0.001, r=0.211) (Figure 5c). Microbiome similarity was positively associated with evolutionary similarity in the fishes for gill (p=0.007, r=0.176), skin (p=0.047, r=0.133), and the hindgut (p=0.001, r=0.306) communities (Figure 5d-f) for Weighted normalized Unifrac. Overall, evolutionary distance in fishes was primarily associated with weighted distances rather than unweighted, suggesting that microbes with higher abundances or compositions had a stronger prediction of host genetic similarity. In addition, within the weighted comparisons, hindgut had the strongest association followed by gill and then skin. Thus, we concluded that both host phylogeny along with the environmental signal (e.g. habitat, benthic substrate, water column depth etc.) were factors important for shaping the mucosal microbial communities of fishes.

Since vertebrates seem to have co-evolved to some extent with their hindgut microbiomes, it is likely that strains from genera that contain known probiotics found in fishes would have higher performance in fishes as compared to terrestrial derived probiotics applied to fishes. Therefore, we next explored the extent by which taxa from genera that contain known probiotics (*Bacillus* and *Lactobacillus*), were present across fish body sites. *Bacillus* was found in a higher frequency of fish species across body sites (gill=48.6%, skin=48.3%, midgut=67%, hindgut=36.5%; percent of species with *Bacillus* present) than *Lactobacillus* (gill=16.8%, skin=15.7%, midgut=19.2%, hindgut=5.9%; percent of species with *Lactobacillus* present) (Supplemental Fig. 4a). Both *Bacillus* and *Lactobacillus* were found in the midgut in a higher number of fish species than other body sites. However, *Bacillus* and *Lactobacillus* made up a very small fraction of the overall community (< 5%), thus future work should include enrichment methods in addition to metagenomics to describe these species (Supplemental Fig. 4b).

### Quantifying the role of sea water and marine sediment as microbial sources for fish mucus

We performed an analysis using SourceTracker2 to better understand the role of the environment in shaping the microbiomes of fish mucosal sites. Specifically, we included 108 marine sediment samples and 60 paired sea water samples from a 10 km transect in San Diego including samples from the beach, sandy soft bottoms, rocky reefs, and bay sand/mud (Figure 6a). The sediment and sea water samples were set as replicate sources with all fish samples included as unique sinks. Across all fish mucosal sites, there appeared to be a gradient of importance with beach sand having the lowest overall microbial contribution to fish mucosal sites followed by marine sediment and lastly marine sea water having the highest contribution of microbes. However, the majority of microbes were still of unknown origin (mean: gill 88.95%, skin 79.06%, midgut 77.03%, hindgut 89.93%) (Figure 6b). Next we asked if certain body sites were more likely to have known microbial sources. Fish body sites differed in the proportion of ASVs derived from sea water sources (P<0.0001, KW=39.66, Figure 6c) with midgut generally having the highest amount of sea water microbes followed by skin and then hindgut and gill. Fish body sites also differed in the proportion of ASVs that were derived from marine sediment sources (P<0.0001, KW=23.41, Figure 6d) again with midgut having the most sediment ASVs followed by skin, hindgut, and then gill. We next tested if within a given body site, there was a difference in the proportion of microbes originating from sea water or sediment. For all body sites, microbes originated more from sea water sources than marine sediment (Mann-Whitney U test, gill P=0.0215, skin P=0.0157, midgut P<0.0001, hindgut P=0.0148) (Figure 6e). Despite the significance, there remained a large range of values across fish species, thus we explored if certain life history traits explained when a fish had higher proportions of microbes originating from sea water or sediment. In comparing the continuous variables for each body site, we found that in the gill and midgut samples, microbial biomass was negatively associated with a SW:SED ratio (enrichment of sediment microbes as compared to sea water). The dorsal:TL ratio (indicates acceleration or fast swimming fish) for the gill, skin, and hindgut samples was positively associated with the SW:SED indicating fish that swam fast had more sea water sourced microbes in those body sites. In addition, the gape to TL was positively associated with SW:SED in the skin and hindgut while trophic level was positively associated with SW:SED in hindgut. Mass and condition factor for midgut, was positively associated with SW:SED (Figure 6f). Since body shape morphology with context to swim performance was associated with SW:SED, we next compared if overall body shape as it relates to swimming (metadata column: swim_performance) was also associated. Specifically, we were interested to know if flatfish that have adapted to living in the sand had a higher proportion of microbes originating from sediment. We found significant differences across swim mode (P=0.0158, KW=13.97) and specifically that cruiser/sprinter fish (includes the mackerels, tunas, and jacks, etc) for the skin body site only had a higher SW:SED (median = 7.916) than flow refuging (flatfish and stingrays/skates) (median = −1.203) and manoeuvrer fish (median = −2.794) (Figure 6g).

**Figure 6.**
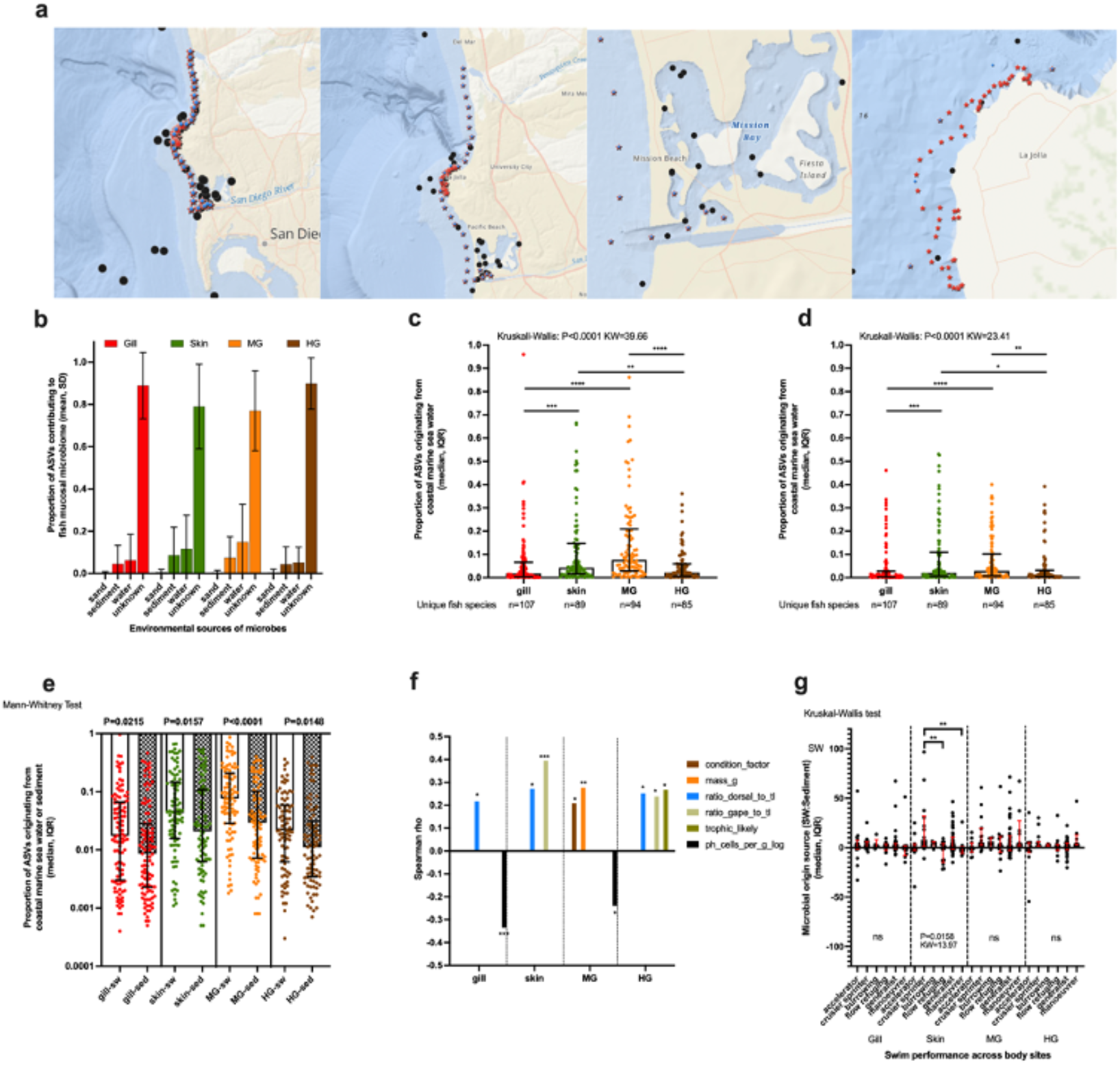
Microbial source tracking analysis. A) Microbial sources of 60 sea water (blue circle) samples taken from 30 unique sampling stations from two time points are distributed on a 10 km transect from Torrey Pines beach to Mission Bay. Microbial sources of 108 marine sediment samples (red stars) from San Diego coastal environment includes 60 paired samples (same locations as sea water) from the same 10 km transect along with 58 samples from the various reef habitats near La Jolla. B) Sourcetracker2 analysis of likely sources for the four body sites of the fish comparing contributions of beach sand, marine sediment, sea water, and ‘unknown’. Unknown refers to microbes from an unknown source which could include diet and other animals or locations not sampled. C) Specific microbial contributions of sea water to the four mucosal body sites and D) specific microbial contributions of marine sediment to the four mucosal body sites (statistical testing across body sites using Kruskal-Wallis, Benjamini-Hochberg multiple comparisons test 0.05 FDR). E) Proportion of microbes likely originating from the sea water vs. sediment for each unique body site (sea water vs. sediment pairwise comparison for each body site using Mann-Whitney test P<0.05). F) Comparison of the ratio of sea water ‘SW’ and marine sediment ‘SED’ against various continuous fish life history metadata variables for each unique body site (Spearman correlation P<0.05). G) Comparison of the SW:SED ratios across the habitats from which the fish live. Comparisons performed on each unique body site (Kruskal-Wallis test, P<0.05).

### Gamma diversity analysis across vertebrates

We performed a meta-analysis focusing on hindgut fecal samples that were the body site of broadest interest in the field and thus had the most samples. Hindgut microbial gamma diversity was compared on rarified data from a total of 569 unique species across the five vertebrate classes (fishes n=73, birds n=216, mammals n=208, reptiles n=52, and amphibians n=20) to test whether total microbial diversity ‘gamma’ was greater in animals with an older evolutionary history (Figure 7a). Mammals had the highest total number of amplified sequence variants ‘ASVs’ (35,105), followed by birds (19,384 ASVs), and then fishes (9,567 ASVs) (Figure 7b). The majority of ASVs were unique to each broad vertebrate class (e.g. mammal, reptile, amphibian, bird, or fish) and not shared with other classes. Of the classes, fishes had the highest percentage of unique ASVs that were not found in other classes (92%), while mammals were second highest at 87% (Figure 7b).

**Figure 7.**
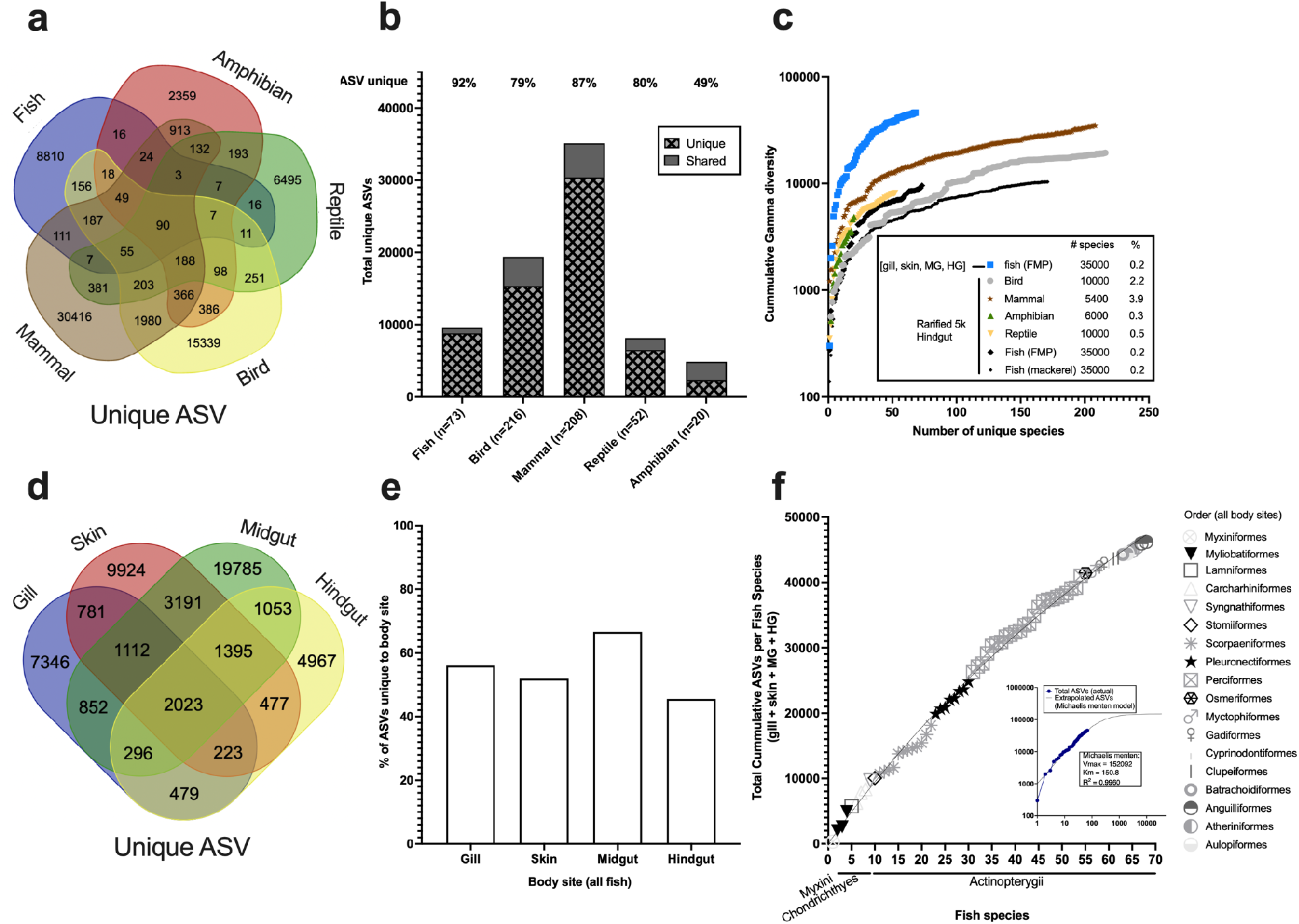
Total microbial diversity across vertebrate hindguts and within multiple body sites of fish. a) Hindgut microbiome samples from 569 species of vertebrates were rarified to 5000 reads and unique or shared ASVs determined for each class. b) The percent of unique ASVs only found in a given class (not shared in other classes) as compared to the total ASVs within that class. c) Rarefaction of cumulative gamma diversity as a function of unique vertebrate species. Included is a single fish species, *S. japonicus*, sampled over three years ‘black dots’ and the unrarified FMP samples which had detectable bacteria in all four body sites (gill, skin, midgut, and hindgut). d) Gamma diversity of 68 fish species across four body sites. e) Percentage of unique ASVs associated with a given body site across the 68 fish species. f) Rarefaction curve of increasing gamma diversity (inclusive of four body sites) as a function of increasing fish species.

Since the total number of species sampled differed across vertebrate classes, we next compared total cumulative microbial diversity on an additive basis for individual species for both the rarified hindgut samples for all vertebrates along with the full unrarified FMP dataset (including other body sites: gill, skin, midgut, and hindgut) of fishes collected from SoCalUSA. When comparing hindgut samples only, at 50 different animal species, mammals had the greatest gamma diversity followed by reptiles and then fishes with birds the lowest. Multiple replicates (50 fish replicates across three years and four seasons) of a single species of fish, *Scomber japonicus*, was included as reference for comparison purposes, although this was similar in magnitude (total unique microbial hindgut diversity at 50 replicates) to birds (Figure 7c).

We next assessed gamma diversity strictly in fish samples that had sufficient microbial sequences in all four body sites (n=68 species) based on the sample exclusion criteria calculated from Katharoseq^11^ (Figure 2). Because we can estimate microbial biomass on a per sample basis based on read counts for the FMP dataset, we included samples as unrarified to enable a better estimate of total diversity. Midgut samples overall had the greatest number of ASVs unique to that body site (19,785 ASVs) followed by skin (9,924 ASVs), and gill (7,346 ASVs) with hindgut (4,967) having the lowest overall gamma diversity (Figure 7d). Next we calculated the extent by which those ASVs made up the total diversity in a given body site (unique ASVs/total ASVs). The midgut also had the greatest proportion of ASVs which were unique to that body site (66.6%) followed by gills (56%), skin (52%), and hindgut (46%) (Figure 7e). We conclude that hindgut samples in fishes are the body site with the lowest total microbial diversity and lowest unique body site associated diversity. Thus, comparisons across vertebrates should begin to include other body sites that may harbor more microbial biomass and diversity.

Lastly, we attempted to assess the feasibility of estimating the total number of V4 region ASVs across all of the estimated 35,000 extant species of fishes. To do this, we performed a rarefaction plot on increasing fish diversity with total cumulative gamma diversity of fishes. Gamma diversity was calculated per fish species by combining all unique ASVs across the gill, skin, MG, and HG. The fish species were arranged on the x-axis by class (with oldest fishes first) and then alphabetically by order and family. Results of the model suggested that saturation occurred after sampling around 1000 species of fish and there was an estimated total of 152092 total unique ASVs associated with fishes (Figure 7f). On the contrary, visual inspection suggests a very small change in the overall trajectory that suggests an accurate estimate of gamma diversity is not possible without many more species, perhaps hundreds or even thousands of additional species. While the goodness of fit for the model is quite high (R2=0.9960), it is entirely possible that we are severely underestimating total diversity, because saturation of the curve is not easily apparent. Also, our study was limited by focusing on marine temperate fishes and few tropical fishes so it was likely underestimating the diversity, especially because it excludes tropical coral reef associated fishes. For future research on gamma diversity across other vertebrates, we recommend comparisons of multiple body sites along with the application of the KatharoSeq^11^ methodology and back calculation as demonstrated in this study to enable the estimation of actual microbial biomass in a given sample.

## DISCUSSION

Here we investigated the ecological and biological drivers of bacterial diversity associated with fish mucosal sites (gill, skin, midgut, and hindgut) across 101 fish species from the EPO along with gill samples from an additional 17 species (15 unique) from the Western Atlantic. We curated a list of both categorical and continuous metadata types to describe the biology of the fish including habitat, diet, and swim performance. In addition we tested the effects of host phylogeny on microbial diversity. Lastly we performed an extensive meta-analysis of gamma diversity of marine fishes as compared to other vertebrate classes and show the importance of including multiple body sites along with microbial biomass in measures of diversity.

Of all the factors tested across the 101 fish species, the ‘body site’ was the strongest predictor of microbial community composition followed by general depth, habitat benthic substrate, and fish microbial biomass. Body site is frequently one of the strongest predictors when comparing single fish species including Atlantic salmon ^14^, Pacific Chub Mackerel ^7,14^, Yellowtail Kingfish ^15^, and Southern Bluefin Tuna ^16^, but this was the first to demonstrate the effect across a range of fish species representing a diverse phylogenetic sampling. This suggests that there are conserved aspects of the body sites across fish lineages, which select for or enrich certain microbial communities. We hypothesize this could be due to body site differences in microbial exposure, immune function, mucus chemistries, morphology, and host anatomy and physiology ^17^. Microbial biomass, habitat depth 1 (shallow ‘neritic’, mid water ‘mesopelagic’, or deep sea ‘bathypelagic or abyssal’), and substrata group (pelagic, soft bottom, rock associated, deep water benthic) were also top predictors of beta diversity across the entire dataset suggesting that habitat had an influence on the microbiome of fishes.

Each body site had specific associations with the various ecological and biological parameters. We observed decreasing microbial diversity in the gills of larger fishes and hypothesize that high performance swimming fishes may have adaptations related to keeping gills clear of microbial fouling to maintain respiratory performance. Fish skin primarily functions as a protective barrier by preventing invasion of pathogens, but in some species the skin can have additional physiological roles such as a source of gas exchange ^18^. Here, skin microbiomes were primarily explained by the type of bottom structure of the habitat from which the fish lives. For shallow environments, the benthic substrate (e.g. mud, sediment, rocky reef) will likely have a stronger contribution of microbes directly to the water column as compared to deeper water systems and therefore may explain how sediment types can influence external microbiomes of fishes ^19,20^. The gastrointestinal tract can vary in morphology and function across fishes with many differing types of stomachs, lengths, and enzyme productions ^21–23^.

Several unique observations with the gill microbiomes in regards to the ecology of fishes suggests a novel evolutionary role. Fishes that were morphologically built for high acceleration (higher dorsal length to total length ratio), had more sea water associated microbes contributing to the gill, skin, and hindgut as compared to sediment microbes. Fishes with a higher microbial biomass in the gills were associated with a higher proportion of sediment sourced microbes as compared to seawater. For neritic fishes, we observed that gill microbial biomass was negatively associated with increasing distance from shore. Combined, we hypothesize that fishes that live closer to shore, such as in the intertidal or subtidal zones as opposed to pelagic fishes, will have lower physiological requirements for swim performance.Therefore, as a potential tradeoff, these fishes will have higher microbial biomass accumulating in the gills as a result of their closer proximity to or in contact with marine sediments. Gills are the source of both gas exchange and nitrogenous waste excretion in fishes and are composed of a generalized conserved morphology with gill arches, filaments, and lamellae. Understanding how microbes may enhance or disrupt these physiological processes will be important areas of research in the future; especially as it relates to aquaculture production of pelagics ^24^.

Contrary to expectations, we did not observe a direct association between trophic level and alpha diversity. We estimated trophic level by two primary methods. First, we estimated the trophic level by using previously documented diet data derived from the literature. In addition, we used the ratio of ‘relative intestinal length: total body length <RIL:TL>” as an indicator of trophic level^25^. Fishes that are more herbivorous will have a higher RIL greater than 1 and upwards of 5-30 whereas carnivorous fishes will generally be much lower < 1 ^26^. Previous work in mammals and fishes has shown that broad trophic levels are generally associated with hindgut microbiome diversity ^12,27,28^. In temperate marine ecosystems, herbivorous marine fishes are rare and therefore it is possible that our limited sampling of the lower trophic extremes could have led to a lack of signal in our study. We did however show that beta diversity between the midgut and hindgut was generally smaller (more similar) in fishes of lower trophic level based on the diet data. This would suggest that microbial differentiation is greater at the start vs. end of the intestine in more carnivorous fishes. Although we did not measure stomach content or relative intestinal content, it is possible that the higher trophic fishes have lower feeding frequencies and thus higher rates of fasting in the wild which has been shown to be a strong predictor of gut microbial communities ^29^. It’s also possible that herbivorous fishes, which feed at a higher frequency rate ^30^, may have overall more similar microbial communities at the proximal and terminal ends of the gut for enhanced nutrient digestion ^31^. Our study did attempt to collect mostly adult sized fish but it’s possible that age could be a confounding variable as herbivorous fish when juveniles are known to have a higher trophic diet ^32^.

Phylosymbiosis occurs when the “microbiomes recapitulate the phylogeny of the host” and is primarily studied in guts of invertebrates and mammals ^33^. Our study showed that the hindgut, gill, and skin microbiomes are more similar in fishes that are more genetically similar. To our knowledge, this is the first study in vertebrates to comprehensively evaluate and show phylosymbiosis, in the context of branching length, occurring across multiple body sites. In addition, the discovery of possible phylosymbiosis occurring in the fish gill has not been shown previously. A positive association between the microbiome and host phylogeny is an important finding for guiding future probiotic discovery as most current probiotics used in aquaculture are derived from terrestrial livestock. For vertebrates, phylosymbiosis has primarily been investigated in mammals with a focus on the ‘internal’ gut microbiome ^33–35^. Only a few studies have investigated phylosymbiosis on animal surfaces such as mammal skin ^36^ (38 species, 10 orders) ^13^, bird feathers (7 species, 1 order) ^37^, Chondrichthyes vs Osteichthyes skin (9 species, 9 orders) ^38^, tropical reef fish skin (44 species, 5 orders) ^39^. It is likely that previous attempts to evaluate phylosymbiosis in fishes have been limited due to limited sampling across evolutionary time scales. In addition, since habitat is an important driver of the fish microbiome, it is important to account for this by having enough samples across diverse habitats as well. For fish, our results suggest that phylosymbiosis is strongest in the hindgut with the gill and skin following in importance in the context of weighted measures. For gut comparisons, future studies should aim to investigate the importance of reproductive strategies in fishes (viviparity vs ovoviviparity vs. oviparity) to determine if phylosymbiosis and potentially co-evolution is stronger in fishes which utilize viviparity. Another important aspect to focus on in future analyses is how microbial diversity corresponds with hosts with high species radiation but shallow overall branching length “low genetic divergence” such as some freshwater cichlids ^40^.

Microbial source tracking analyses showed that sea water contributes more microbes to the fish mucosal environment as compared to sediment. Across body sites, midgut overall had the most microbial sources identified whereas the hindgut and gill had the least (most unknowns). For all body sites however, the majority of microbes were of unknown origin which suggests that further research needs to be conducted to establish a holistic microbial library of the entire marine ecosystem from this California Current Ecosystem region. This sampling effort should include sampling of sediments from deeper depths, sea water from bays and offshore environments, representatives from the thousands of marine invertebrate species, and hundreds of macroalgae species.

Understanding the factors that shape the microbial ecology across vertebrates and specifically fishes remains challenging. Our study showed that amongst vertebrates, fishes have the most unique assemblage of total microbial diversity, which we hypothesized corresponds to the diverse evolutionary history and habitat types exhibited by fishes. For gut samples, 92% of ASVs found in fishes were not found in other classes such as mammals, birds, reptiles, and amphibians despite mammals and reptiles having higher gamma diversity at a 50 species cross-section. Some caveats of our design however was that although all of our fish samples were wild, many of the mammal, bird, reptile, and amphibian samples were from zoo collections ^4^. Zoo and wild samples may differ in diversity due to restrictions in diet but also in feeding frequency ^41^. In addition, the actual sampling of organisms was not random across the tree but instead was opportunistic based on available data, thus future studies should revisit these comparisons when higher species representation is obtained especially from wild samples across a large home range. We demonstrated across one of the largest samplings of marine fishes species to date, that mucosal diversity is greater in body sites outside of the hindgut, particularly the midgut, gill, and skin. This is in contrast to mammals and specifically humans that seem to have the highest proportion of microbial diversity concentrated in the hindgut ‘stool’^42^. Few studies look at the foregut of mammals compared to the hindgut making it difficult to speculate how selection may differ across vertebrate classes. For fish however, because of this drastic difference in midgut to hindgut diversity, it’s possible that the majority of midgut microbes are simply from diet and ingested seawater, representing a reservoir of microbes for intestinal colonization. Our study suggested that mammals may have expanded diversity in the hindgut as compared to other vertebrates, including fishes. Whereas fishes, may have higher cumulative diversity spread across other body sites. However, one major caveat to gamma diversity comparisons in vertebrates is the morphological differences in body sites. Fishes and to some extent amphibians have gills whereas mammals, birds, and reptiles have lungs for breathing. External surfaces such as the ‘skin’ also differ widely. The ideal comparison would be to process and extract the entire animal at once, but typically this isn’t feasible due to size limitations. In order to do broad gamma diversity comparisons across vertebrates, one may need to focus on conserved body sites including reproductive organs, oral cavity, anus with the understanding that one would be excluding diversity elsewhere. Higher hindgut diversity in mammals could be explained by the higher occurrence of herbivory in mammals ^43^. Our study did not include tropical marine fishes from coral reef ecosystems as we concentrated on the California Current marine ecosystem ‘CCE’. The CCE does have tropical fish, but they are associated with rocky reef benthic habitat. The addition of gut microbiomes from coral reef fishes, which are primarily localized in tropical environments, may influence this outcome as herbivory is higher in that habitat for fishes ^44^. In mammals, one of the most important drivers of hindgut microbial diversity is the complexity and physiology of the gut, namely if hindgut fermentation occurs ^45^. The adaptation of herbivory may have led to expanded physiological and morphological attributes in the foregut and hindgut leading to a novel ecological niche for microbial colonization and symbiosis of fermentative taxa ^5^.

Because few vertebrate microbiome studies are inclusive of multiple body sites, it is difficult to compare cumulative gamma diversity of the fishes here to other vertebrates aside from our observation of higher microbial diversity in external body sites such as the gill and skin in fishes. Our attempts to estimate total microbial diversity across body sites extrapolated to 35,000 fish species demonstrates that despite an incredibly rich dataset with a large range of fish diversity, our investigations of microbial diversity in fishes are superficial. We suspect the expanded diversity in external sites in fishes may be explained by an evolutionary pressure of a high exposure rate to microbes in the aquatic environment. This may have led to the diversification of the immune system including mucosal site specific lymphoid associated tissue and mucus production ^46–48^. Microbiome diversity in the hindgut but not external sites is partially associated with immune system complexity of the hosts ^49^. A follow up study would be to compare the immune components (e.g. gene expression, protein, or metabolome profiles) across these different body sites (gill, skin, gut) within an individual fish to determine the extent the host immune system influences (permits or excludes) microbial diversity at a local body site level. In addition, fishes that are naturally exposed to higher microbial diversity in their life history (oligotrophic vs eutrophic or pelagic vs. benthic) may exhibit differences in the evolution of their immune system.

### Conclusion

To fully measure and evaluate microbial diversity in vertebrates, our study showed it is imperative to design studies that include microbiome samples from a broad phylogenetic sampling of hosts along with multiple body sites. In addition, we demonstrated the importance of including microbial biomass measurements in the context of diversity estimates. Our method to interpolate and approximate microbial biomass of fish mucosal microbiome sites from standards is an important improvement to the field that should be applicable across future datasets.

## METHODS

### Fish microbiome project (FMP) metadata

#### Sampling

A total of 101 fish species were collected from Southern California, primarily in San Diego County, USA ranging from latitude (31.435833 to 33.142589) and longitude (−118.20833 to −117.20879). Fishes were collected using a variety of methods but primarily hook and line, spear, or trawls (for deep sea fishes). All samples with the exception of the thresher shark and seven gill shark were immediately stored on dry ice and then kept in the −80C until dissection. The mentioned sharks were dissected immediately and body sites frozen on dry ice followed by storage in −80C freezer ^50^. Four body sites were processed for microbiome analyses including the gill, skin, midgut and hindgut. For gill samples, in most instances, the whole gill (left 2nd filament) was used. When gills were too large, three slivers of the entire filament were collected from the top, middle and bottom of the gill arch. Skin samples were primarily from scraped mucus. Midgut and hindgut were digesta material either posterior of the stomach at the beginning of the GI (midgut) or at the anus (hindgut). Details of all fishes used in the study along with its corresponding metadata can be found in (Supp file FMP.metadata) with the corresponding details of that metadata in the data dictionary (Supp file FMP.datadict). Taxonomy assignments were done using fishtreeoflife.org and fishbase.org, and ultimately NCBI taxonomy ^51^.

#### Trophic assignment

Diet preferences of each species was determined through literature searches including secondary literature databases such as fishbase.org. Where diets differed for juvenile and adult stages, all components were included in the metadata. A coarse estimate of trophic level was determined based on the general diet. If a fish rarely eats fish or if fish are a minor component of the diet, they are still considered in the trophic numbering where 1=herbivore (consumes primary producers), 2= primary carnivore (consumes primary consumers such as zooplankton or other invertebrates), 3= secondary carnivore (consumes small fishes), 4= tertiary carnivore (consumes large carnivorous fishes), or 5= quaternary carnivore (consumes high trophic fish or marine mammals). In addition to assigning the trophic level based on the literature (trophic_likely), we can also estimate the trophic level based on the ratio of GI length to Total length. A high GI:TL (>2 or 3) will be associated with more herbivorous fishes whereas a short ratio (<1) is associated with carnivorous fishes or a higher trophic level.

#### Reproduction

Fish are classified in their reproduction method (oviparous = external egg fertilization, ovoviviparous = internal fertilization nourished by egg yolk with live birth, viviparous = internal fertilization nourished by mother gas exchange with live birth) in the metadata column (Reproductive_process).

#### Habitat

Multiple classification methods were used to capture the habitat diversity. The actual minimum and maximum depths are included as (depth_low and depth_high). From here, a broad classification of either shallow, midwater, or deep was used to indicate fishes dwelling between (0-200, 200-1000 m, and <1000 m) (metadata column=habitat_depth_level1). In the next level (metadata column=habitat_depth_level2), we segregate shallow by either intertidal (0-~10 m) or neritic (0-200 m). Fishes dwelling primarily between 200-1000 m are indicated as mesopelagic. Fishes which dwell between 200-1000 m yet are largely demersal are further classified as mesopelagic/benthopelagic. Fishes living between 1000 - 4000 m are labeled bathypelagic and lastly if dwelling < 4000 m are abyssopelagic. Note, the classifications related to habitat depth are based on where the fish primarily reside rather than at what depth they were caught as most have large ranges for vertical migration throughout the day. For instance, most of the deep sea fishes (classified as bathypelagic) were caught at 500 m. As for salinity tolerance (metadata column = salinity_tolerance) fishes which can live in estuaries are labeled as ‘brackish’ and all others as ‘marine’. Since some fishes undergo greater migrations, we have added a column to indicate (salinity_tolerance_migrations). Fish are either marine (if only in ocean), oceanodromous (migrate long distances in ocean), brackish (spend part of all of life in brackish or estuarine waters), anadromous, and catadromous. The ocean basin from which fish were caught is indicated by (ocean_basin).

#### Swim performance

Swim acceleration, swim endurance, and the ‘dorsal length to total length (TL)’ ratio are all measures of swim performance. For swim acceleration and endurance, we assigned values to each fish based on the body shape morphometrics previously described for the fish ^52^. For acceleration and endurance we assigned either a 1, 2, or 3 with the acceleration value of 3 indicative of high speed (e.g. a barracuda) whereas a high endurance value (3) is indicative of high endurance. In addition, we can describe the capacity for fast swimming based on the placement of the dorsal fin on the fish body. A more forward dorsal fin will be associated with slower swimming fishes whereas fast swimming fishes will generally have a dorsal fin more towards the tail. The dorsal length to TL ratio is a morphometric measure with a higher ratio being indicative of fast swimming or ability to accelerate quickly.

### Microbiome Processing

#### Fish microbiome project samples

The fish microbiome project data is held in Qiita study ID 13414 ^53^. For each extraction set of 96 samples, a set of 8 positive controls were included. These positives were from either the Zymo mock community (Zymo Cat# D6300) or single microbial isolates with known cell concentration based on plate counts. Each isolate or mock was then processed by a serial dilution to extinction to get a range of biomass as input following the Katharoseq protocol ^11,54^. Following the Earth Microbiome Project protocols ^3^, all DNA extractions were processed using the Qiagen PowerMag kit with a modification in that the lysis step was performed in single 2 ml tubes while the cleanup performed on the KingFisher Flex robots using the magnetic bead cleanup. This modification was done to reduce well to well contamination which is common when doing plate based lysis using vortexing for DNA extractions ^55^. In addition, fish species were randomly assigned across the plate so that groups of similar phylogeny were not necessarily clustered together on the platemap but instead scattered. All gDNA was eluted to 60 ul. For the PCR step, a total of 2 ul of DNA was used in a miniaturized 10 ul PCR reaction ^56^. After PCR, an equal volume of DNA was pooled from all libraries into a single sequencing pool following the Katharoseq protocol. Equal volume pooling is essential to enable downstream quantification across libraries since sequencing read counts correlate to original DNA input ^1111,54^ to the PCR and subsequently cell biomass to extraction. The final sequencing library was then cleaned up using 1x AMPure beads to remove PCR contaminants. Samples were run on three separate sequencing instruments including: MiSeq Nano 2×250 bp (artifact ID 102012), NovaSeq 2×250 bp (Lane 1 artifact ID 112123, Lane 2 artifact ID 112121), and a MiSeq 2×150 bp (artifact ID 113069). All samples were processed according to the Earth Microbiome through Qiita ^53^, trimmed to 150 bp and ASVs generated using Deblur v1.1.0 and only ASVs passing the positive filter step were used ^57^. The analysis artifact ID on Qiita is 46238.

#### Microbial biomass estimation

We developed a methodology to enable the estimation of microbial biomass from a sample. First you process the standards through the standard Katharoseq pipeline to determine the limit of detection ^11^. Briefly, this requires one to have standards (microbial isolates or mock communities) with known cell concentrations which can be determined using standard plate counts or other methods. A (8, 10-fold) serial dilution of the standards is performed, and then extracted alongside actual samples, with a minimum of 32 total positives included on an experiment (4 replicates per dilution). It is critical that all samples and standards are processed identically (same elution volume, same volume into PCR, etc). It is also critical that the actual biomass of the samples of interest are determined (weighed out). After sequencing, one uses the relationship between read counts and known cell counts of the standards to determine the limit of detection (typically at a setting of 0.9). One excludes all samples and controls with reads lower than this cutoff. This is described in great detail in the original Katharoseq publication. To estimate the biomass of samples, one then log 10 transforms the read counts and cell counts of the positive controls which pass the LoD. The relationship is modeled using linear regression and the slope and y intercept is then used to estimate the “cell counts” of the actual samples with the log 10 read counts used as input in a similar methodology employed by qPCR. One then must account for the amount of DNA volume used in the library prep along with the elution volume used in DNA extraction. Finally, one must also normalize based on the original biomass used in the extraction to get a final estimate of the microbial cells per gram of “e.g. fish tissue”. This method with all normalization steps is now available as a qiime2 plugin “katharoseq”. https://github.com/biocore/q2-katharoseq.

### Microbiome Analyses

Alpha, beta, and gamma diversity measures are quantitative measures used in ecology to assess biological diversity as a general function of some discrete area ^58^. These values were generally calculated in Qiita using Qiime 2 ^59^. For alpha diversity, we calculated richness (total unique observed ASVs), Shannon, and Faith’s Phylogenetic Diversity. For testing categorical variables against alpha diversity measures, we use the nonparametric Kruskal-Wallis test with a Benjamini Hochberg FDR of 0.05. When comparing continuous metadata factors to alpha diversity, we use Spearman correlation. For beta diversity, we used Unweighted Unifrac which focuses on rare taxa along with Weighted normalized Unifrac which is more heavily weighted or influenced by abundant taxa ^60^. Statistical testing of metadata categories was performed using PERMANOVA with 999 permutations and a significance of 0.001^61^. Gamma diversity was defined by the total sum of diversity in a given unit which could be a class of vertebrates (e.g. mammals, fish, etc.) or within a given species (gill + skin + gut, etc.).

#### Meta-analysis of gamma diversity across vertebrate classes

We first performed a meta analysis comparing hindgut microbiome diversity across numerous vertebrate species. Specifically, we utilized data from previously published or available datasets (Qiita Study ID - Artifact ID - European Nucleotide Archive accession: 11721 - 111895 [ERP109537] ‘mackerel 1 year’; 13066 - 87276 ‘mackerel year 2-3’; 10353 - 59141 [ERP106745] ‘Malawi manure’; 13414 - 102012, 112121, 112123, 113069 ‘FMP’; 12227 - 67067, 67063 [ERP120036] ‘Australia fish’; 11166 [ERP118494] - 56540, 82398, 82409, 82395, 82512, 82400, 82965; 11687 - 85793, 58423 ‘SD coastal microbiome’; 12769 - 81577 ‘SD map and bioreactor’ ^4,7,62^. Only hindgut samples from fish were used initially and only a single replicate per species was used to eliminate pseudoreplication as a confounding factor. All samples were rarefied to 5000 reads to ensure an even comparison of gamma diversity. A total of 73 fish species (out of ~35,000), 216 bird species (out of ~10000), 208 mammal species (out of ~5400), 52 reptile species (out of ~10000), and 20 amphibian species (out of ~6000) were included in the analysis representing a total of ~0.86 % of the total vertebrate diversity (569 species out of 66,400 total vertebrate species). The frequency table used to generate this analysis (Supplemental Source 1. Gamma.freqtable.allspecies.txt). The total list of ASVs were tabulated for each class and then visualized using a venn diagram (Figure 7a) using http://bioinformatics.psb.ugent.be/webtools/Venn/. For each vertebrate class (fish, mammal, bird, reptile, and amphibian), the total unique ASVs were calculated by taking the total number of ASVs only found in a given class and dividing by the total number of ASVs found within that class. The total ASVs of a class include ASVs shared amongst other classes (Figure 7b). An example for fish would be (8,810 = total unique ASVs only found in fish / 9,567 = total ASVs found in fish = 92.1%). Cumulative gamma diversity as a function of sequencing additional species was tabulated for all classes specifically for hindgut microbiomes (Figure 7c). In addition, samples from the fish microbiome project ‘FMP’ which has successful sequencing at each of the four body sites were also included as a comparison (n=68 species) (Figure 1). The total unique microbiome diversity across all four body sites were used as the gamma diversity metric in this case with additive unique ASVs calculated for increasing species sequenced. All additional metrics of gamma diversity for the FMP samples (Figure 7d-f) were calculated using the 68 fish species which had successful sequencing results for all four body sites.

Cross vertebrate fecal analysis (Lactobacillus discovery) [Qiita analysis 46249] Studies included: 10353 (malawi manure), FMP, VMP 11166, Salmon/SBT/YTK studies 12227, HMP 1927).

#### Fish Microbiome Project analysis

For the FMP samples, samples with less than 1150 reads were excluded leaving a total of 373 successful samples. For the unrarified raw table, this included 55069 ASVs which included 1165 chloroplast ASVs which were then removed. For the rarified table (1150 reads), a total of 22605 ASVs passed filter including 562 chloroplast associated ASVs which were then removed.

#### Probiotic discovery

The majority of probiotics used in aquaculture are derived from terrestrial sources. Potential bacterial probiotics included any ASV within the following genera: *Bacillus, Lactobacillus, Enterococcus*, and *Carnobacterium* (others are *Lactococcus*, and *Weissella*) as those have been shown to have a beneficial role as immunostimulants or growth promoters ^63^. For our analysis, we focused on *Bacillus* and *Lactobacillus* as they are the most commonly used in industrial applications. We identified the prevalence and estimated the relative abundance of these two genera across the FMP dataset.

#### Phylogenetic analysis

Phylosymbiosis can be generally estimated by comparing the phylogenetic distances of the host (in this case fish) to the microbiome similarities of these hosts ^33^. To compare the effect of host phylogeny on the microbiome, we generated a tree of all of the fish species used in this dataset using timetree.org ^64^. We then used the estimates of evolutionary distance (divergence time) for each pairwise comparison of hosts and tested for associations between host divergence time and gut microbiome divergence using Mantel tests (*mantel.rtest* function of the R ade4 package). We performed this test for each body site uniquely and determined significance at P<0.05. For the microbiome similarity metrics, we included Unweighted Unifrac and Generalized Weighted Unifrac distances.

## Data availability

All samples will be uploaded to EBI and all analyses will be made public upon acceptance and/or during peer review. All source data used in figures along with code for alpha diversity and phylosymbiosis analyses can be found at: https://github.com/jminich444/Fish_Microbiome_Project

The FMP101 analysis was generated from Qiita analysis ID: 46251 “FMP 2021-07-14_v3 deblur”. This includes the following artifacts: 102012, 112121, 112123, and 113069 from Qiita study ID 13414.

The probiotic analysis where Lactobacillus and Bacillus ASVs were tabulated across vertebrates can be found in Qiita analysis ID 46249 “FMP: broad gamma vertebrate analysis”.

Sourectracker2 analysis used biom tables generated from Qiita analysis ID: 49005 “FMP101_SDcoastal_[ST2]”. This included the following studies: <Qiita study ID 12769; artifact ID 132396> “San Diego Coastal Microbiome Map” and <Qiita study ID 13414; artifact IDs 135520, 135536, 135719> “Fish Microbiome Project”.

The gamma diversity analysis of comparing the rarified hindgut samples across 569 vertebrates can be found in Qiita analysis ID 46287 “FMP_ultra metaanalysis”. This same analysis ID also contains the analysis of the unrarified fish gamma diversity.

The code and methods for the biomass estimation can be found and used as a Qiime2 plugin called ‘katharoseq’. https://github.com/biocore/q2-katharoseq

## ACKNOWLEDGEMENTS

We would like to thank the California Department of Fish and Game for permitting this project (2016 Scientific Collecting permit DFW 1379: DocID: D-0018712881-8). We thank Mike Shane and Brice Semmens along with their corresponding organizations for enabling us to collect opportunistic samples which would have otherwise been discarded such as the Hubbs- SeaWorld Research Institute and the California Cooperative Oceanic Fisheries Investigations. We thank the crew of the R/V Robert Gordon Sproul including Phil Zerofski. We especially thank all of the fishers who donated their fish for analysis.

## AUTHOR CONTRIBUTIONS

JJM designed the experiment, collected samples, processed samples, curated metadata, analyzed data, and wrote and revised the manuscript. AH performed the phylogenetic (phylosymbiosis) analysis and helped write and revised the manuscript. JV helped process samples and helped curate metadata, specifically the swim mode data. BWF and ZS helped in collecting samples and identifying fish species along with reviewing and revising the manuscript. EM helped in sample collection and sample processing along with reviewing and revising the manuscript. MS helped in sample collection. EEA, RK, and TPM assisted in oversight, mentoring, writing, and revising the manuscript.

