## Supplemental_figures for "Fish microbiomes 101: disentangling the rules governing marine fish mucosal microbiomes across 101 species"

| Class | Order | Family | Species/FMP number | |
|---|---|---|---|---|
| Actinopterygii | Albuliformes | Albulidae (bonefishes) | Albula vulpes<br>bonefish<br>FMP 50 | 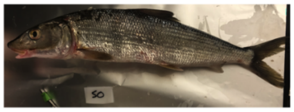 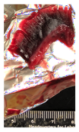 |
| | Anguilliformes | Anguillidae (freshwater eels) | Anguilla rostrata<br>american eel<br>FMP 113 (A12) | 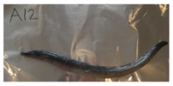 |
| | | Muraenidae (moray eels) | Gymnothorax mordax<br>california moray<br>FMP 68 | 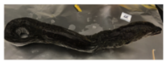 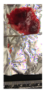 |
| | | Nemichthyidae (snipe eels) | Nemichthys scolopaceus<br>snipe eel<br>FMP 78 | 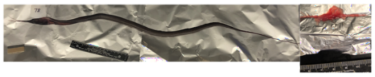 |
| | | Nettastomatidae (duckbill eels) | Facciolella equatorialis<br>dog face witch eel<br>FMP 86 | 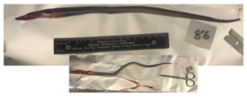 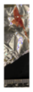 |
| | Argentiniformes | Bathylagidae (deep-sea smelts) | Leuroglossus stilbius<br>CA Smoothtongue<br>FMP 77 | 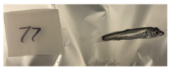 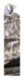 |
| | | | Lipolagus ochotensis<br>Earned Blacksmelt<br>FMP 70 | 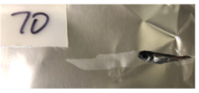 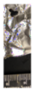 |
| | Atheriniformes | Atherinopsidae (neotropical silversides) | Atherinops affinis<br>topsmelt<br>FMP 99 | 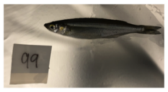 |
| | | | Atherinopsis californiensis<br>jacksmelt<br>FMP 19 | 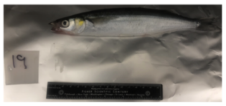 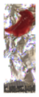 |
| | Aulopiformes | Synodontidae (lizardfishes) | Synodus lucioceps<br>CA lizardfish<br>FMP 51 | 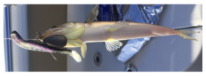 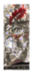 |
| | Batrachoidiformes | Batrachoididae (toadfishes) | Opsanus tau<br>oyster toadfish<br>FMP 112 (A11) | 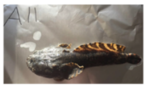 |
| | | | Porichthys myriaster<br>specklefin midshipman<br>FMP 48 | 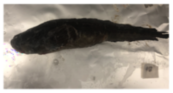 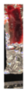 |
| | | | Porichthys notatus<br>plainfin midshipman<br>FMP 72 | 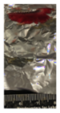 |
| | Beloniformes | Belonidae (needlefishes) | Strongylura marina<br>Atlantic needlefish<br>FMP 111 (A10) | |

| Class | Order | Family | Species/FMP number | |
|---|---|---|---|---|
| Actinopterygii | Beryciformes | Melamphaidae (bigscale fishes/ Ridgeheads | Scopelogadus bispinosus<br>Twospine Bigscale<br>FMP 87 | 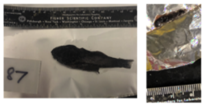  |
| | Blenniformes | Blenniidae (combtooth blennies) | Chasmodes bosquianus<br>Striped Blenny<br>FMP 110 (A9) | 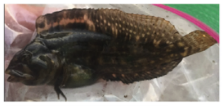 |
| | | | Hypsoblennius gilberti<br>Rockpool blenny<br>FMP 34 | 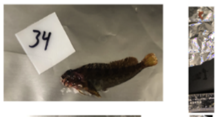  |
| | | Clinidae (clinids) | Gibbonsia elegans<br>Spotted Kelpfish<br>FMP 33 | 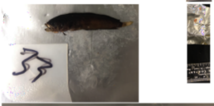  |
| | | | Heterostichus rostratus<br>Giant Kelpfish<br>FMP 23 | 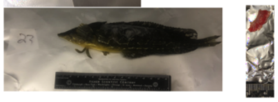 |
| | Carangiformes | Carangidae (jacks and pompanos) | Caranx crysos<br>Blue Runner<br>FMP 114 (A13) | 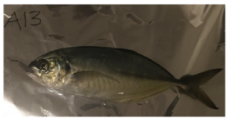  |
| | | | Seriola lalandi<br>CA yellowtail<br>FMP 79 | 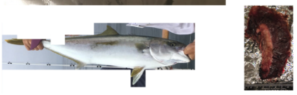 |
| | | | Trachurus symmetricus<br>Pacific jack mackerel<br>FMP 20 | 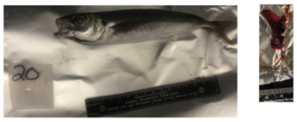  |
| | | Coryphaenidae (dolphinfishes) | Coryphaena hippurus<br>Dolphinfish<br>FMP 115 (A14) | 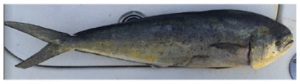 |
| | | | Coryphaena hippurus<br>Dolphinfish<br>FMP 97 | 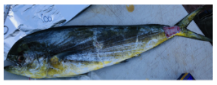 |
| | | Sphyraenidae (barracudas) | Sphyraena argentea<br>Pacific 'CA' Barracuda<br>FMP 11 | 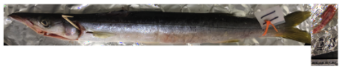 |
| | Centrarchiformes | Girellidae | Girella nigricans<br>Opaleye<br>FMP 22 | 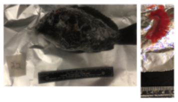 |
| | | Kyphosidae (sea chub) | Kyphosus azurea<br>Zebra Perch<br>FMP 21 | 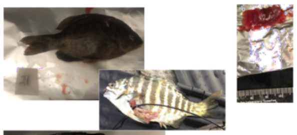 |
| | | | Medialuna californiensis<br>Halfmoon<br>FMP 54 | 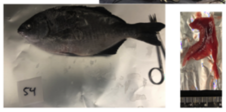 |

| Class | Order | Family | Species/FMP number | |
|---|---|---|---|---|
| Actinopterygii | Clupeiformes | Clupeidae<br>(herring, shad,<br>sardine, &<br>menhadens) | Brevoortia tyrannus<br>Atl menhaden<br>FMP 109 (A8) | 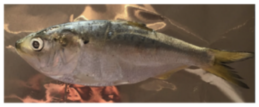 |
| | | | Sardinops sagax<br>sardine<br>FMP 60 | 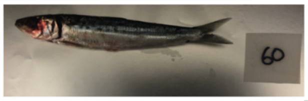 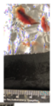 |
| | | Engraulidae<br>(anchovies) | Engraulis mordax<br>CA anchovy<br>FMP 40 | 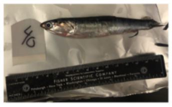 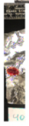 |
| | Cyprinodontiformes | Fundulidae<br>(topminnows<br>and killisfishes) | Fundulus heteroclitus<br>mummichog<br>FMP 108 (A5) | 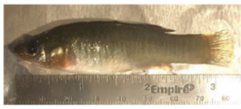 |
| | | | Fundulus parvipinnis<br>California killifish<br>FMP 96 | 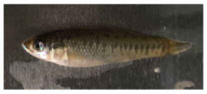 |
| | | | Lucania parva<br>Rainwater Killifish<br>FMP 116 (A18) | |
| | Gadiformes | Macrouridae<br>(grenadiers<br>or rattails) | Nezumia stelgidolepis<br>California Grenadier<br>FMP 93 | 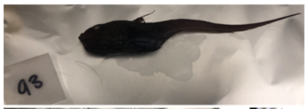 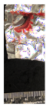 |
| | | Merlucciidae<br>(merluccid hakes) | Merluccius productus<br>North Pacific Hake<br>FMP 74 | 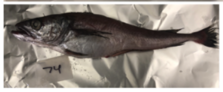 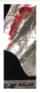 |
| | | Moridae<br>(morid cods) | Physiculus rastrelliger<br>Hundred Fathom Mora<br>FMP 64 | 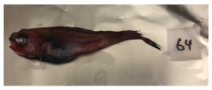 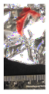 |
| | Gobbiformes | Gobiidae<br>(gobbies) | Clevelandia ios<br>Arrow Goby<br>FMP 95 | 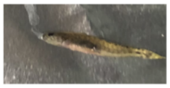 |
| | | | Gillichthys mirabilis<br>Longjaw Mudsucker<br>FMP 100 | 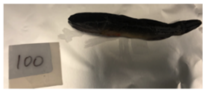 |
| | Labriformes | Labridae<br>(wrasses) | Halichoeres semicinctus<br>Rock Wrasse<br>FMP 8 | 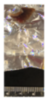 |
| | | | Oxyjulis californica<br>Senorita<br>FMP 24 | 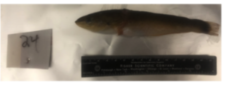 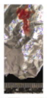 |
| | | | Semicossyphus pulcher<br>CA Sheephead<br>FMP 18 | 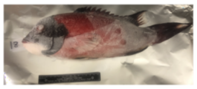 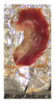 |

| Class | Order | Family | Species/FMP number | |
|---|---|---|---|---|
| Actinopterygii | Myctophiformes | Myctophidae (lanternfishes) | Ceratoscopelus townsendi<br>Dogtooth Lampfish<br>FMP 91 | 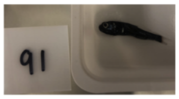  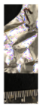 |
| | | | Nannobrachium ritteri<br>Broadfin lampfish<br>FMP 88 | 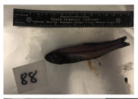  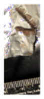 |
| | | | Triphoturus mexicanus<br>Mexican Lampfish<br>FMP 61 | 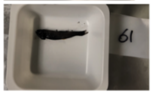  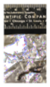 |
| | Pempheriformes | Polyprionidae (wreckfishes) | Stereolepis gigas<br>Giant Seabass<br>FMP 92 | 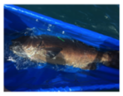 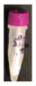 |
| | Perciformes | Agonidae (poachers) | Xeneretmus latifrons<br>Blacktip Poacher<br>FMP 69 | 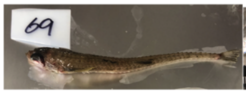  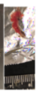 |
| | | Cottidae (sculpins) | Leptocottus armatus<br>Pacific Staghorn Sculpin<br>FMP 16 | 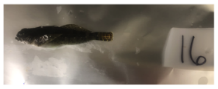  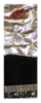 |
| | | Embiotocidae (surfperches) | Embiotoca jacksoni<br>Black Perch<br>FMP 9 | 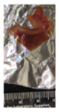 |
| | | | Phanerodon furcatus<br>White Seaperch<br>FMP 7 | 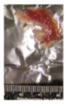 |
| | | | Brachyistius frenatus<br>Kelp Perch<br>FMP 53 | 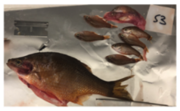  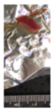 |
| | | | Rhacochilus toxotes<br>Rubberlip Seaperch<br>FMP 52 | 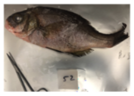 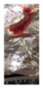 |
| | | Haemulidae (grunts) | Anisotremus davidsonii<br>Xantic Sargo<br>FMP 10 | 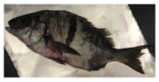 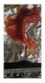 |
| | | | Xenistius californiensis<br>CA Salema<br>FMP 45 | 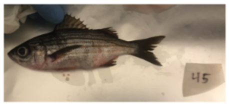  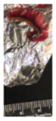 |
| | | Malacanthidae (tilefishes) | Caulolatilus princeps<br>Ocean Whitefish<br>FMP 2 | 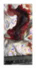 |

| Class | Order | Family | Species/FMP number | |
|---|---|---|---|---|
| Actinopterygii | Perciformes | Moronidae<br>(temperate<br>basses) | Morone saxatilis<br>Striped Bass<br>FMP 120 (A23) | 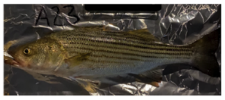 |
| | | Pomacentridae<br>(damselfishes) | Chromis punctipinnis<br>Blacksmith<br>FMP 36 | 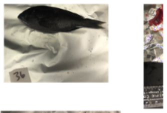 |
| | | Sciaenidae<br>(drums &<br>croakers) | Bairdiella chrysoura<br>Silver Perch<br>FMP 107 (A4) | 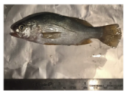 |
| | | | Cheilotrema saturnum<br>Black Croaker<br>FMP 102 | 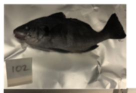 |
| | | | Cynoscion parvipinnis<br>Shortfin Weakfish<br>FMP 49 | 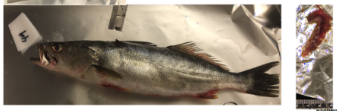 |
| | | | Menticirrhus undulatus<br>CA Kingcroaker<br>FMP 30 | 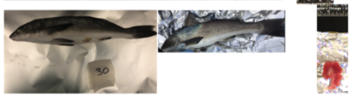 |
| | | | Roncador stearnsii<br>Spotfin Croaker<br>FMP 46 | 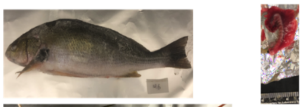 |
| | | | Seriphus politus<br>Queen Croaker<br>FMP 13 | 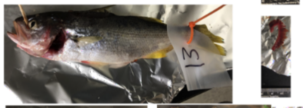 |
| | | | Umbrina roncador<br>Yellowfin Drum<br>FMP 44 | 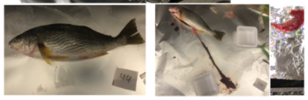 |
| | | Scorpaenidae<br>(scorpionfish) | Scorpaena guttata<br>California Scorpionfish<br>FMP 41 | 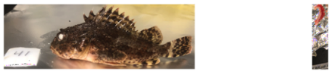 |
| | | Sebastidae<br>(rockfish) | Sebastes auriculatus<br>Brown Rockfish<br>FMP 42 | 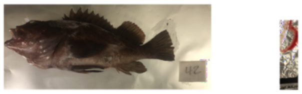 |
| | | | Sebastes carnatus<br>Gopher Rockfish<br>FMP 5 | 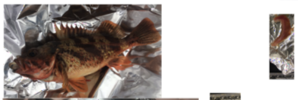 |
| | | | Sebastes chlorostictus<br>Greenspotted Rockfish<br>FMP 39 | 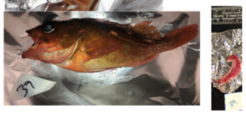 |
| | | | Sebastes constellatus<br>Starry Rockfish<br>FMP 4 | 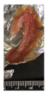 |
| | | | Sebastes dallii<br>Calico Rockfish<br>FMP 27 | 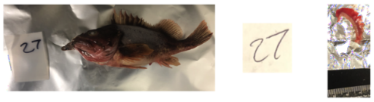 |

| Class | Order | Family | Species/FMP number | |
|---|---|---|---|---|
| Actinopterygii | Perciformes | Sebastidae (rockfish) | Sebastes diploproa<br>Splitnose Rockfish<br>FMP 63 | 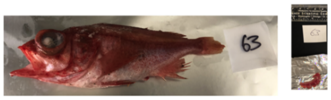 |
| | | | Sebastes hopkinsi<br>Squarespot Rockfish<br>FMP 38 | 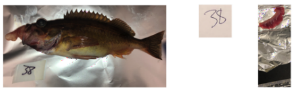 |
| | | | Sebastes miniatus<br>Vermillion Rockfish<br>FMP 28 | 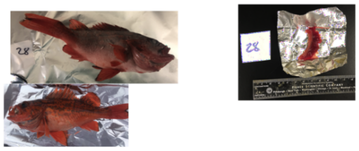 |
| | | | Sebastes mystinus<br>Blue Rockfish<br>FMP 6 | 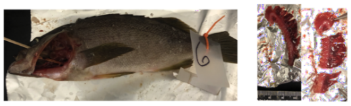 |
| | | | Sebastes semicinctus<br>Halfbanded Rockfish<br>FMP 29 | 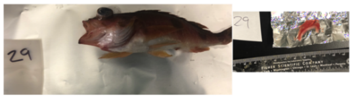 |
| | | | Sebastes serriceps<br>Treefish<br>FMP 17 | 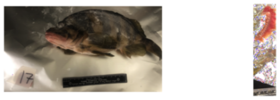 |
| | | | Sebastes umbrosus<br>Honeycomb Rockfish<br>FMP 3 | 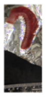 |
| | | Serranidae (sea basses, groupers, & fairy basslets | Centropristis striata<br>Black Seabass<br>FMP 106 (A3) | 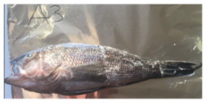 |
| | | | Paralabrax clathratus<br>Kelp Bass<br>FMP 1 | 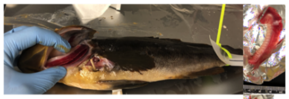 |
| | | | Paralabrax maculatofasciatus<br>Spotted Sand Bass<br>FMP 35 | 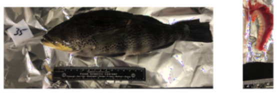 |
| | | | Paralabrax nebulifer<br>Barred Sand Bass<br>FMP 12 | 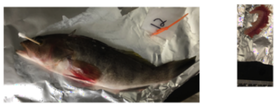 |
| | | Triglidae (searobins) | Prionotus carolinus<br>Northern Searobin<br>FMP 105 (A2) | 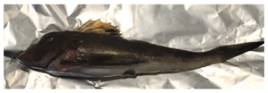 |

| Class | Order | Family | Species/FMP number | |
|---|---|---|---|---|
| Actinopterygii | Perciformes | Zoarcidae<br>(eelpouts) | Lycodes cortezianus<br>Bigfin eelpout<br>FMP 75 | 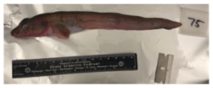 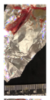 |
| | | | Lycodes diapterus<br>Black Eelpout<br>FMP 82 | 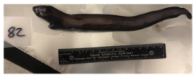 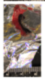 |
| | | | Lycodes pacificus<br>Blackbelly Eelpout<br>FMP 85 | 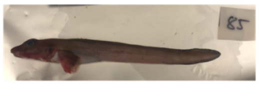 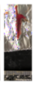 |
| | | | Lyconema barbatum<br>Bearded Eelpout<br>FMP 76 | 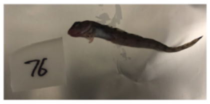 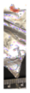 |
| | Pleuronectiformes | Cynoglossidae<br>(tonguefish) | Symphurus atricaudus<br>California Tonguefish<br>FMP 43 | 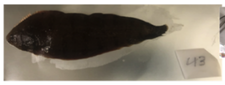 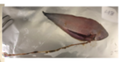 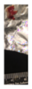 |
| | | Paralichthyidae<br>(large tooth<br>flounders) | Citharichthys sordidus<br>Pacific Sanddab<br>FMP 71 | 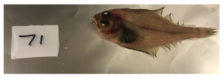 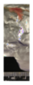 |
| | | | Citharichthys xanthostigma<br>Longfin Sanddab<br>FMP 26 | 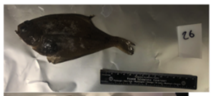 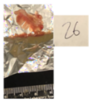 |
| | | | Paralichthys californicus<br>California Flounder (Halibut)<br>FMP 37 | 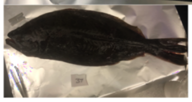 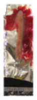 |
| | | | Paralichthys dentatus<br>Summer Flounder<br>FMP 104 (A1) | |
| | | Pleuronectidae<br>(righteye<br>flounders) | Glyptocephalus zachirus<br>Rex Sole<br>FMP 66 | 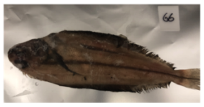 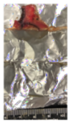 |
| | | | Hypsopsetta guttulata<br>Diamond Turbot<br>FMP 101 | 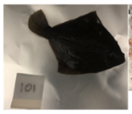 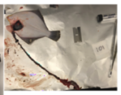 |
| | | | Lyopsetta exilis<br>Slender sole<br>FMP 62 | 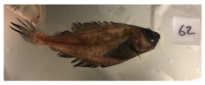 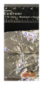 |
| | | | Microstomus pacificus<br>Dover Sole<br>FMP 65 | 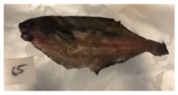 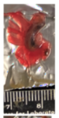 |
| | | | Parophrys vetulus<br>English Sole<br>FMP 73 | 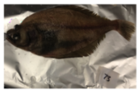 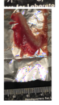 |

| Class | Order | Family | Species/FMP number | |
|---|---|---|---|---|
| Actinopterygii | Scombriformes | Pomatomidae (bluefishes) | Pomatomus saltatrix<br>Bluefish<br>FMP 119 (A22) | 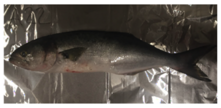 |
| | | Scombridae (mackerel, tunas, bonitos) | Katsuwonus pelamis<br>Skipjack Tuna<br>FMP 98 | 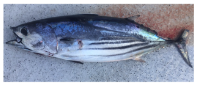 |
| | | | Sarda Chiliensis<br>Pacific Bonito<br>FMP 58 | 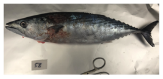 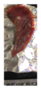 |
| | | | Scomber japonicus<br>Pacific Chub Mackerel<br>FMP 103 | 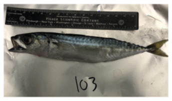 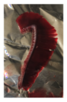 |
| | | | Thunnus albacares<br>Yellowfin Tuna<br>FMP 118 (A21) | 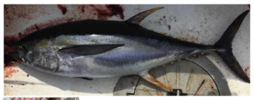 |
| | | | Thunnus albacares<br>Yellowfin Tuna<br>FMP 94 | 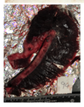 |
| | Stomiatiformes | Sternoptychidae (marine hatchetfish) | Argyropelecus affinis<br>Pacific Hatchetfish<br>FMP 90 | 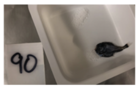 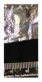 |
| | | | Sternoptyx pseudobscura<br>Highlight Hatchetfish<br>FMP 89 | 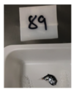 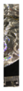 |
| | | Stomiidae (barbeled dragonfishes) | Stomias atriventer<br>Black Belly Dragonfish<br>FMP 83 | 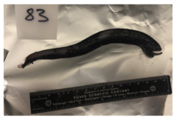 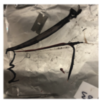 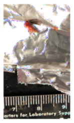 |
| | Syngnathiformes | Syngnathidae (pipefishes & seahorses) | Syngnathus leptorhynchus<br>Bay Pipefish<br>FMP 25 | 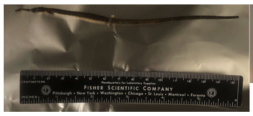 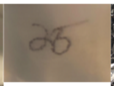 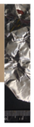 |

| Class | Order | Family | Species/FMP number | |
|---|---|---|---|---|
| Chondricthyes | Carcharhiniformes | Scyliorhinidae (catsharks) | Apristurus brunneus brown catshark FMP 84 | 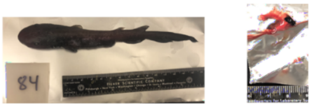 |
| | | Triakidae (houndsharks) | Mustelus californicus gray smooth-hound FMP 14 | 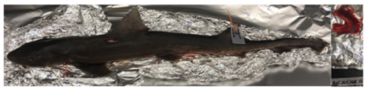 |
| | | | Triakis semifasciata leopard shark FMP 15 | 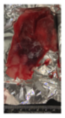 |
| | Heterodontiformes | Heterodontidae (bullhead, horn, or port jackson sharks) | Heterodontus francisci horn shark FMP 32 | 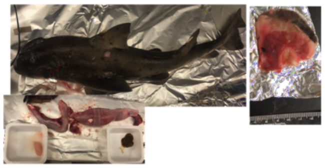 |
| | Hexanchiformes | Hexanchidae (cow sharks) | Notorhynchus cepedianus seven gill shark FMP 55 | 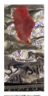 |
| | Lamniformes | Alopiidae (thresher sharks) | Alopias vulpinus thresher shark FMP 59 | 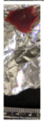 |
| Chondrichthyes | Myliobatiformes | Dasyatidae (stingrays) | Pteroplatytrygon violacea pelagic ray FMP 80 | 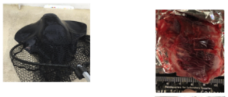 |
| | | Gymnuridae (butterfly rays) | Gymnura marmorata butterfly ray FMP 67 | 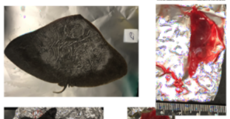 |
| | | Myliobatidae (eagle & manta rays) | Myliobatis californica bat ray FMP 31 | 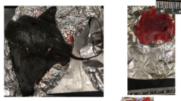 |
| | | Urotrygonidae (American round stingrays) | Urolophus halleri hallers round ray FMP 47 | 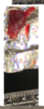 |
| | Rajiformes | Rajidae (skates) | Leucoraja erinacea little skate FMP 117 (A20) | 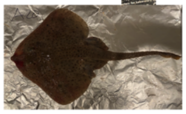 |
| Myxini | Myxiniformes | Myxinidae (hagfishes) | Eptatretus stoutii Pacific hagfish FMP 81 | 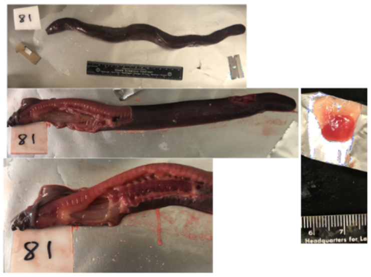 |

Supplemental Figure 1. Atlas of fishes (alphabetical order) used in the fish microbiome project. Black font = fishes from Eastern Pacific; Red font = fishes collected from Western Atlantic.

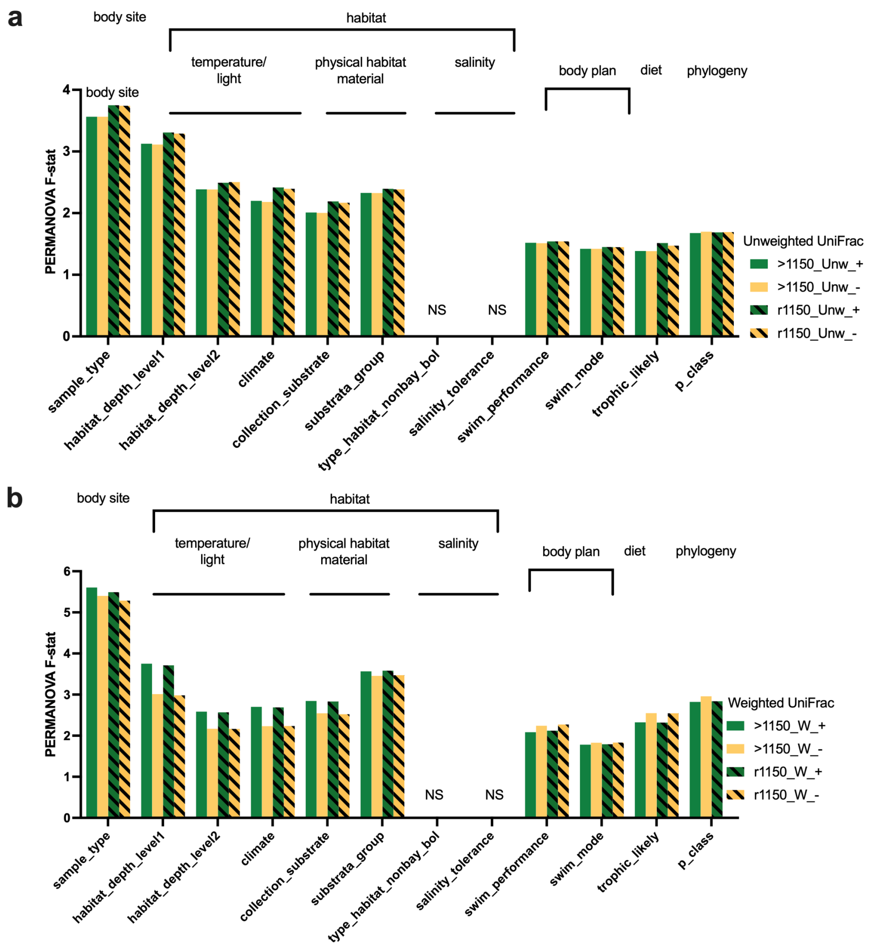


Supplemental Figure 2. Comparison of data processing methods (rarefying and removal of chloroplasts) on beta diversity significance testing using (a) Unweighted Unifrac and (b) Weighted UniFrac distances. Comparison of exclusion of samples with less than 1150 reads (non-rarified) (hashed bars) vs. rarefying at 1150 reads (clear bar). Comparison of datasets with (green bar +) and without (yellow bar -) chloroplast ASVs.

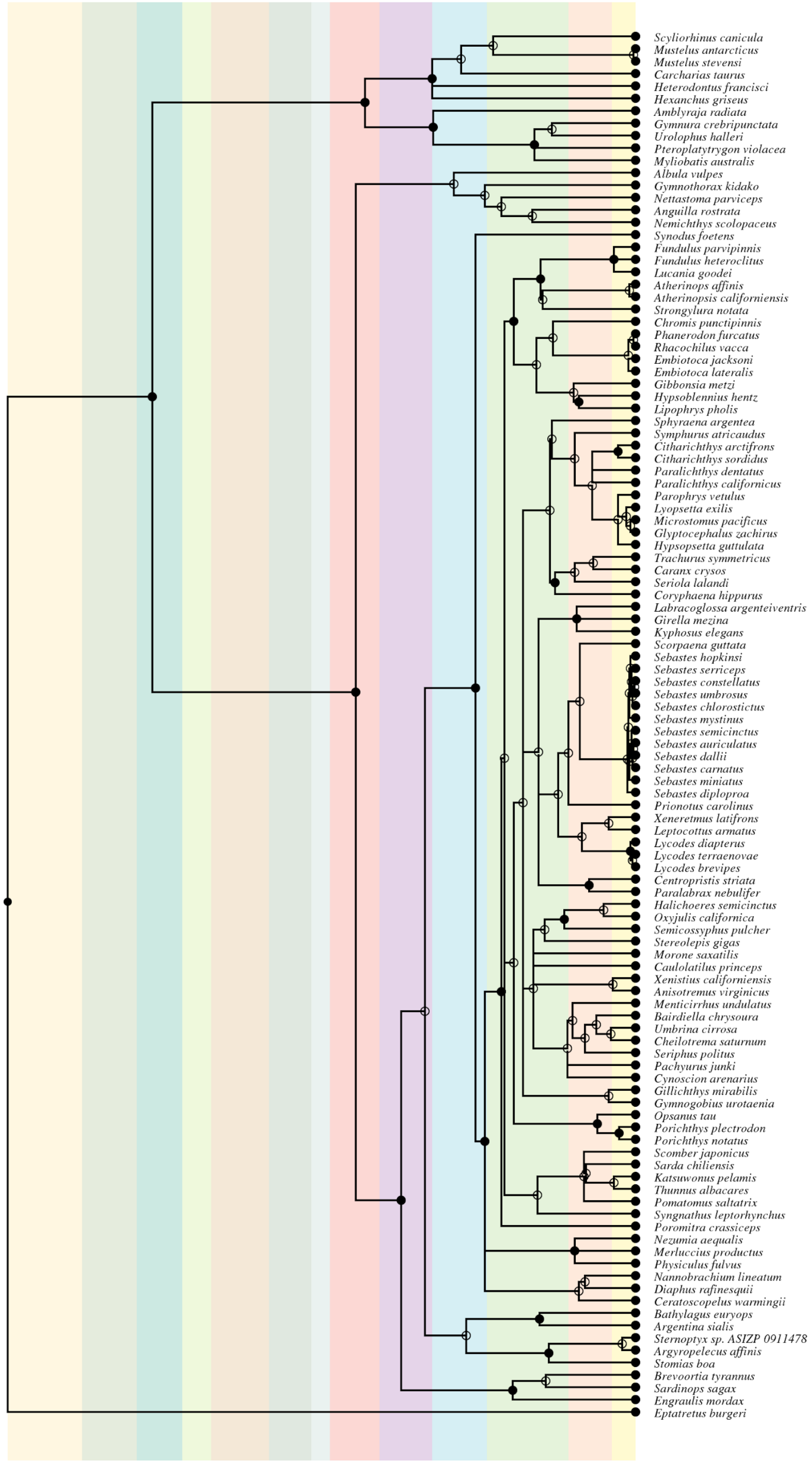


**Supplemental Figure 3. Phylogenetic tree of fish species used in “FMP” beta diversity analysis**

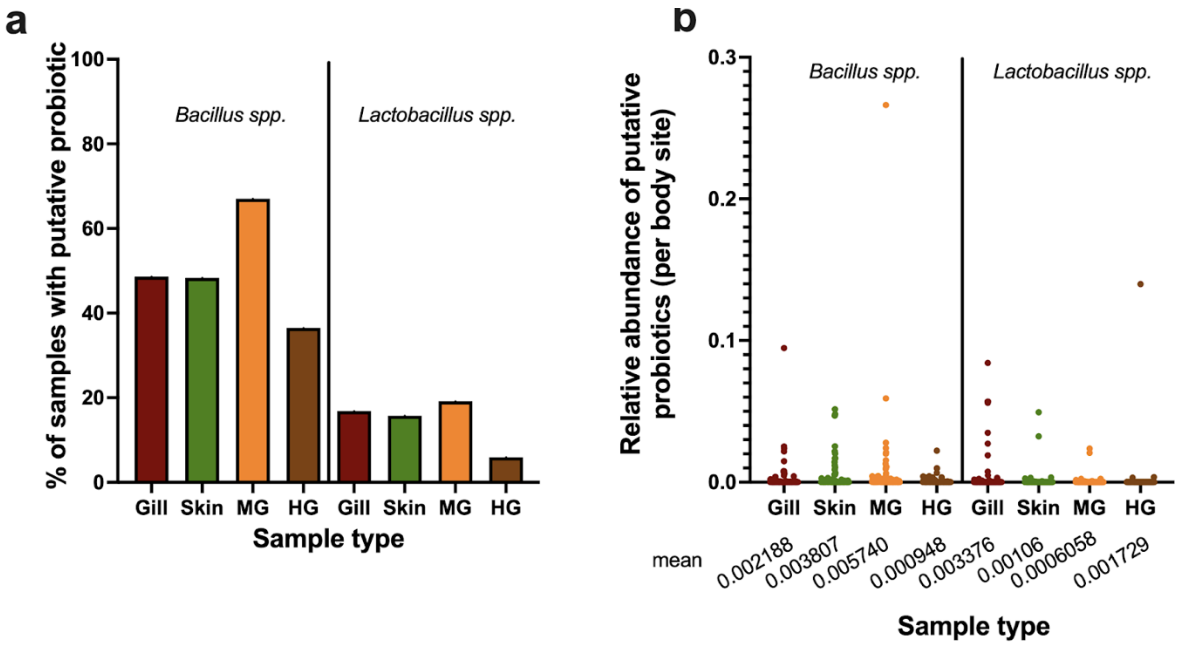


**Supplemental Figure 4 Probiotic. Distribution of putative probiotic ASVs grouped by genera: *Bacillus* or *Lactobacillus.*** a) The total percentage of samples within a given body site having either a Bacillus ASV or Lactobacillus ASV. b) The relative abundances of Bacillus and Lactobacillus ASVs within each body site (including samples with 0 counts).
